# Cells of the human intestinal tract mapped across space and time

**DOI:** 10.1101/2021.04.07.438755

**Authors:** R Elmentaite, N Kumasaka, HW King, K Roberts, M Dabrowska, S Pritchard, L Bolt, SF Vieira, L Mamanova, N Huang, I Goh Kai’En, E Stephenson, J Engelbert, RA Botting, A Fleming, E Dann, SN Lisgo, M Katan, S Leonard, TRW Oliver, CE Hook, K Nayak, F Perrone, LS Campos, C Dominguez-Conde, K Polanski, S Van Dongen, M Patel, MD Morgan, JC Marioni, OA Bayraktar, KB Meyer, M Zilbauer, H Uhlig, MR Clatworthy, KT Mahbubani, K Saeb Parsy, M Haniffa, KR James, SA Teichmann

**Affiliations:** Wellcome Sanger Institute, Wellcome Genome Campus, Hinxton, Cambridge CB10 ISA, UK; Centre for Immunobiology, Blizard Institute, Queen Mary University of London, London E1 2AT, UK; Biosciences Institute, Faculty of Medical Sciences, Newcastle University, Newcastle upon Tyne NE2 4HH, UK; Molecular Immunity Unit, Department of Medicine, University of Cambridge, MRC Laboratory of Molecular Biology, Cambridge, CB2 0QH, UK; European Molecular Biology Laboratory, European Bioinformatics Institute, Wellcome Genome Campus, Cambridge, CB10 1SD, UK; Cancer Research UK Cambridge Institute, University of Cambridge, Cambridge, UK; Structural and Molecular Biology, Division of Biosciences, University College London WC1E 6BT, UK; Department of Histopathology, Cambridge University Hospitals NHS Foundation Trust, Hills Road, Cambridge CB2 0QQ, United Kingdom of Great Britain and Northern Ireland; Wellcome Trust – MRC Cambridge Stem Cell Institute, Anne McLaren Laboratory, University of Cambridge, Cambridge CB2 0SZ, UK; Department of Paediatrics, University of Cambridge, Cambridge CB2 0QQ, UK; Department of Paediatric Gastroenterology, Hepatology and Nutrition, Cambridge University Hospitals Trust, Cambridge, CB2 0QQ, UK; Translational Gastroenterology Unit, John Radcliffe Hospital, University of Oxford, Oxford, OX3 9DU, UK; Department of Paediatrics, University of Oxford, Oxford, OX1 2JD, UK; NIHR Oxford Biomedical Research Centre, Oxford, OX3 7JX, UK; Department of Surgery, University of Cambridge and NIHR Cambridge Biomedical Research Centre, Cambridge, CB2 0QQ, UK; Theory of Condensed Matter Group, Cavendish Laboratory/Department of Physics, University of Cambridge, Cambridge CB3 0HE, UK

## Abstract

The cellular landscape of the human intestinal tract is dynamic throughout life, developing *in utero* and changing in response to functional requirements and environmental exposures. To comprehensively map cell lineages in the healthy developing, pediatric and adult human gut from ten distinct anatomical regions, as well as draining lymph nodes, we used singlecell RNA-seq and VDJ analysis of roughly one third of a million cells. This reveals the presence of BEST4+ absorptive cells throughout the human intestinal tract, demonstrating the existence of this cell type beyond the colon for the first time. Furthermore, we implicate IgG sensing as a novel function of intestinal tuft cells, and link these cells to the pathogenesis of inflammatory bowel disease. We define novel glial and neuronal cell populations in the developing enteric nervous system, and predict cell-type specific expression of Hirschsprung’s disease-associated genes. Finally, using a systems approach, we identify key cell players across multiple cell lineages driving secondary lymphoid tissue formation in early human development. We show that these programs are adopted in inflammatory bowel disease to recruit and retain immune cells at the site of inflammation. These data provide an unprecedented catalogue of intestinal cells, and new insights into cellular programs in development, homeostasis and disease.

## Introduction

Intestinal tract physiology relies on the integrated contribution of epithelial, mesenchymal, endothelial, mesothelial, innate and adaptive immune, and neuronal cell lineages, whose relative abundance and cell networking fluctuate from embryonic development to adulthood. Factors contributing to these dynamics include gut function, environmental challenges and disease states that vary at different life stages. Adding further complexity is that the intestinal tract is formed of distinct anatomical regions that develop at different rates and carry out diverse roles in digestion, nutrient absorption, metabolism, and immune regulation in adulthood.

Single-cell analyses of intestinal organoids from iPSCs and human primary tissue have provided a unique opportunity to study the earliest stages of gut development. Although focused mainly on epithelial cells, they have led to identification of factors driving cellular differentiation ^1,2^ and novel markers of rare cell types ^3^. Analysis of rare fetal intestinal tissues have further resolved the formation of villi-crypt structures and seeding of immune cells into the gut environment ^4,5^.

Similarly, our understanding of the cellular landscape of the adult gut is benefiting from single-cell technologies. We have previously reported regional differences in immune cell activation in the healthy human colon linked to variability in the microbiome composition ^6^ and expression of ligand-receptor pairs between cell populations has been used to infer cellular communication involving enteric neuronal cells subtypes ^7^. Studies comparing inflammatory bowel disease (IBD) samples to healthy controls or non-inflamed tissue have allowed for the identification of a new stromal subtype that expands in disease ^8,9^, clonal expansion of tissue-resident CD8 T cells in disease ^10–12^ and correlation between cellular response and clinical treatment ^13^. While extensive work has been carried out to profile the intestinal tract at single-cell resolution, a holistic analysis of the gut through space (anatomical location) and time (lifespan) is lacking.

Here, we create a broad and deep single-cell census of the healthy human gut, encompassing around 350,000 cells from 11 distinct anatomical sites during embryonic and fetal development, childhood and adulthood. In doing so, we reveal new insights into the epithelial compartment: we identify the presence of BEST4+ enterocytes in both small and large intestine and throughout life, and define a role for IgG sensing by tuft cells in IBD. In the developing enteric nervous system, we identify multiple novel populations, and highlight cell communication networks with stromal cells *via* receptors and ligands associated with Hirschsprung’s disease. In addition, we identify key cells and signaling networks initiating lymphoid structure formation in early human development. Interestingly, the same developmental programmes are adopted to drive recruitment of immune cells during pediatric Crohn’s disease. Thus our high-resolution genomic definition of human cells and gene expression programmes across epithelial, enteric nervous system and immune compartments uncovers new insights into both rare and common diseases of the intestines.

## Results

### Integrated view of human gut throughout life

To investigate cellular profiles and communication networks across the intestinal tract, we performed scRNA-seq on distinct tissue regions of the second-trimester (12-17 postconception weeks (PCW)) fetal and adult (29-69 years) intestine and draining mesenteric lymph nodes (mLN) (Figure 1A & Supplementary Fig. 1A). In adult samples, immune (CD45+) and non-immune (CD45-) cells were separated and loaded for scRNA-seq in equivalent proportions. To represent the gut at embryonic and early postnatal life, we incorporated our previously published embryonic (6-10 PCW; small intestine, large intestine) and pediatric (4-12 years; ileal) scRNA-seq data ^4^.

**Figure 1:**
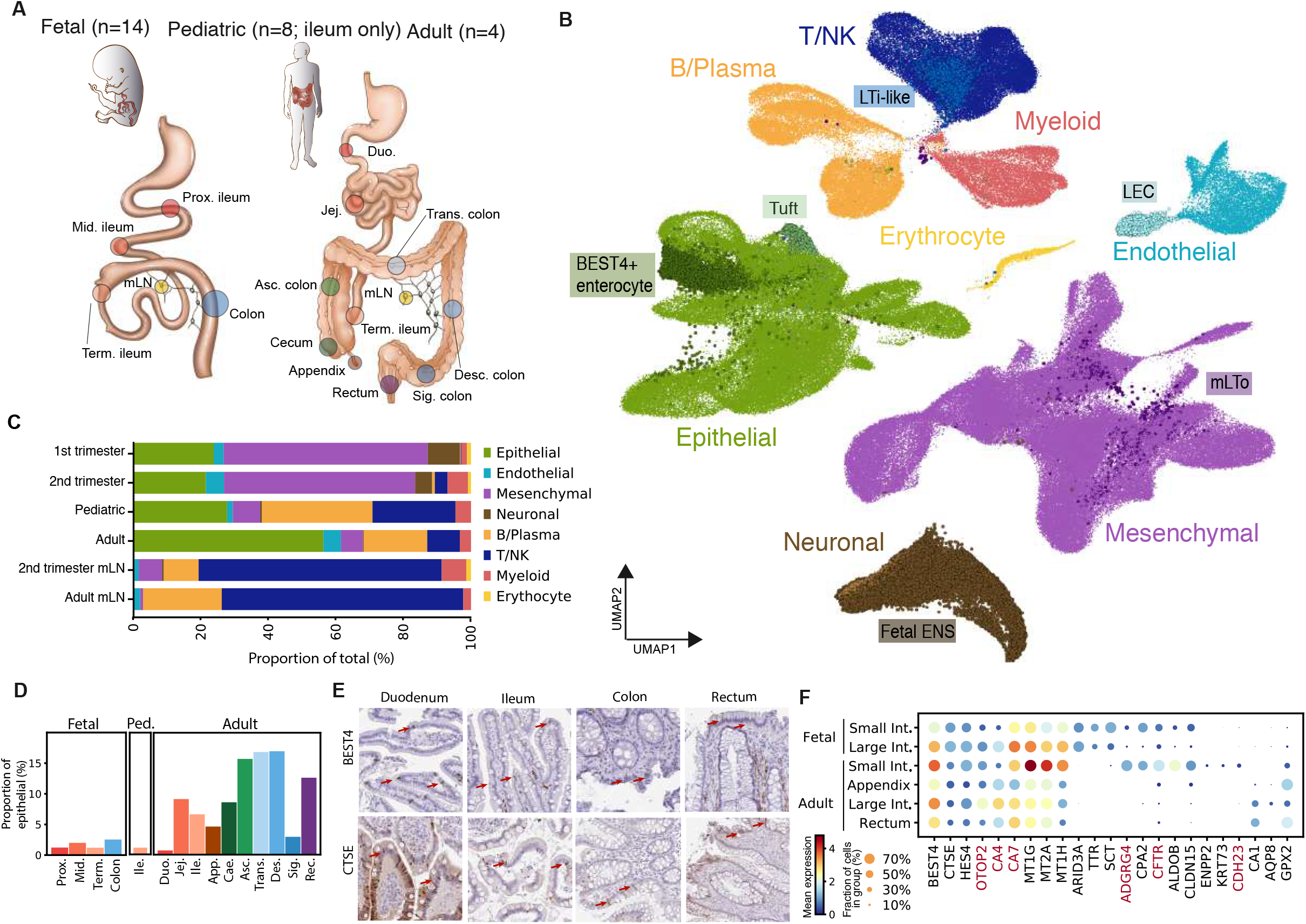
Intestinal cellular census throughout life. A) Schematic of tissue sampling of healthy human gut. Number of individual donors sampled are shown in parentheses. B) UMAP visualisation of cellular landscape of the human intestinal tract coloured by cellular lineage. C) Relative proportions of cell lineage as in B per anatomical tissue. D) Proportions of BEST4+ enterocytes among all epithelial cells per tissue region and developmental stage. E) Expression of BEST4 (antibody: HPA058564) and CSTE (antibody: CAB032687) in gut histological sections from Human Protein Atlas (proteinatlas.org). F) Dotplot with relative expression of small and large common and small intestine specific BEST4+ enterocyte marker genes.

After quality control and doublet removal, the combined dataset comprised over 346,000 intestinal cells (Supplementary Fig. 1B). Leiden clustering and marker gene analysis revealed major clusters of epithelial, mesenchymal, endothelial, lymphocytes, neuronal, myeloid and erythroid cells (Fig. 1B). Fetal gut samples were enriched for mesenchymal cells and relatively high numbers of neuronal cells, with increased abundance of immune cell types from the second trimester onwards (Figure 1C & Supplementary Fig. 2A). Mesenteric lymph nodes, collected from second-trimester onwards, predominantly contained immune cells (Figure 1C & Supplementary Fig. 2A). Further sub-clustering of the cellular populations allowed for identification of 103 cell types and states with specific transcriptional identities (Supplementary Fig. 1B & 2B).

Notably, BEST4+ enterocytes, recently described in the human colon and predicted to transport metal ions and salts ^9,14^, were identifiable in all regions of the fetal and adult intestinal tract suggesting a requirement for these functions in small and large intestines throughout life (Figure 1D-E). Adult small intestinal BEST4+ cells formed a distinct cluster, separated from adult large intestinal and fetal BEST4+ cells (Supplementary Fig. 2C). Using a novel differential cell-type abundance analysis method called “Milo” (Methods), we identified expression signatures specific to small intestinal BEST4+ cells, including adhesion G protein-coupled receptor *ADGRG4* and cadherin *CDH23* (Figure 1F, Supplementary Fig. 2D-E). BEST4+ enterocytes of the adult large intestines expressed higher levels of *OTOP2* (Figure 1F), suggesting a greater propensity for proton transport. While small and large intestine shared expression of the carbonic anhydrase *CA7, CA4* was more highly expressed in the large intestines. Interestingly, small intestinal BEST4+ cells were marked by high expression of chloride channel and cystic fibrosis gene *CFTR* (Figure 1F), which we also observed at the protein level (Supplementary Fig. 2F). Over 60% of cystic fibrosis patients experience intestinal symptoms characterised by inflammation and sticky mucus ^15^. This analysis elucidates distinctions between subtypes of BEST4+ epithelial cells found along the intestinal tract, and specifically implicates small intestinal BEST4+ cells in cystic fibrosis pathogenesis.

Together, this represents an in-depth cell census of the gut and a map of cellular diversity across regions and developmental time. This dataset enables us to determine previously unappreciated patterning of gene expression in the BEST4+ enterocyte compartment along the intestine.

### Diversity of epithelial cells in the human intestinal tract

Next, we further investigated the composition of the prenatal and postnatal gut epithelium in detail. This lineage is formed of major classes of absorptive cells, secretory and enteroendocrine cells (EECs) (Figure 2A, Supplementary Fig. 2B & 3A). The absorptive cells separate into clusters of pediatric and adult small intestinal enterocytes, and large intestinal colonocytes in the pediatric and adult samples. The secretory cells include goblet, tuft, paneth and microfold (M) cells, as well as the precursor states.

**Figure 2:**
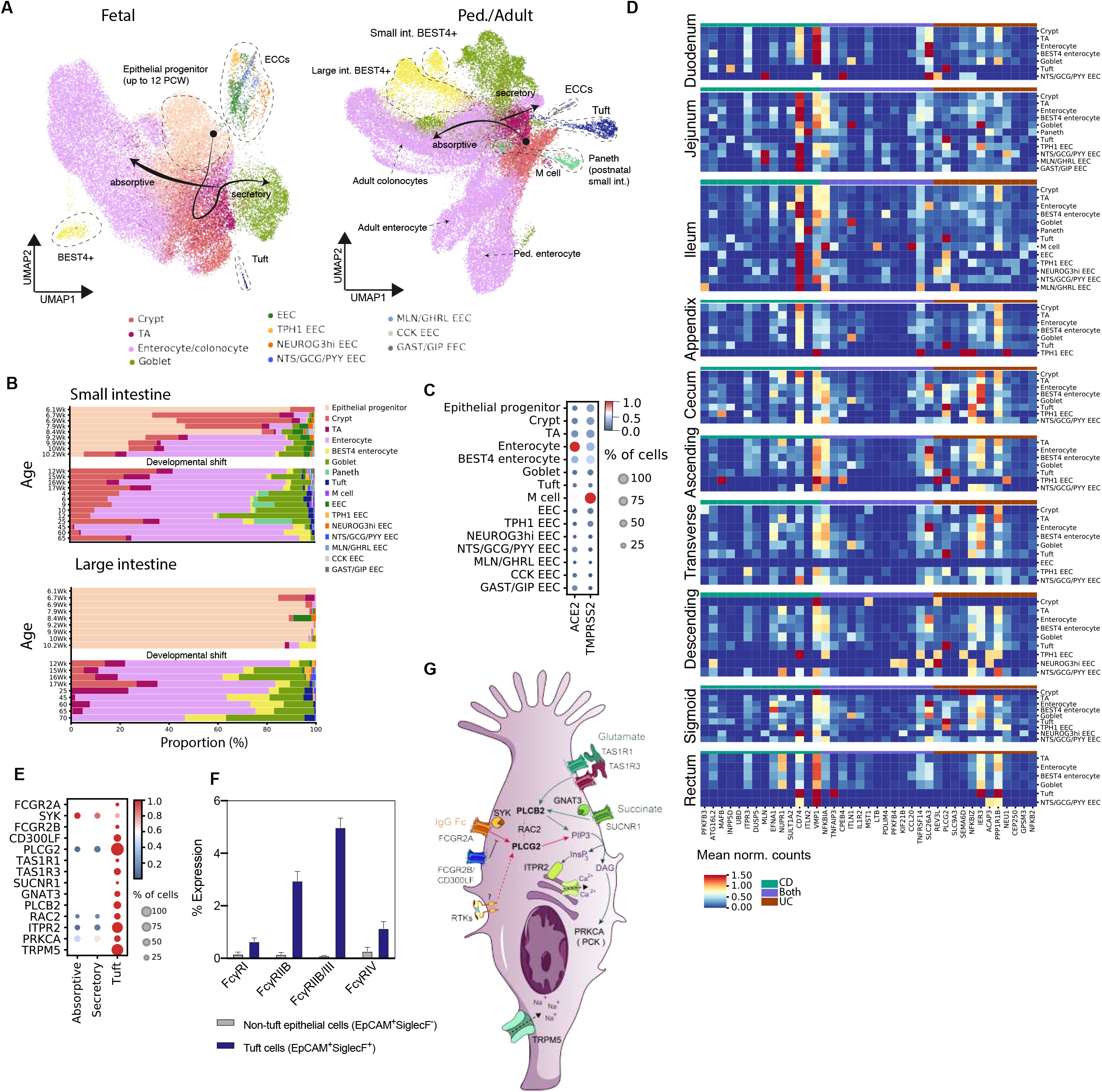
Identification of FCGR2A signalling in tuft cell and its contribution to IBD. A) UMAP visualisation of fetal (left) and postnatal (pediatric and adult; right) epithelial cells coloured by cell type. Key cell types discussed within text are circled with a dashed line. B) Relative proportions of subtypes within total epithelial cell faction separated by donor age (row). Units of age is years unless specified as weeks. C) Dot plot of expression of *TMPRSS2* and *ACE2* by epithelial cell types in the fetal intestine. Here and in later figures, color represents maximum-normalized mean expression of marker genes in each cell group, and size indicates the proportion of cells expressing marker genes. D) Heatmap displaying the expression of Crohn’s disease and ulcerative colitis genes from GWAS by pediatric and adult gut epithelial cells. Only differentially expressed genes between epithelial cells types and with a mean expression of 0.5 normalised counts are included. Scale bar has an upper limit of 1.5 normalised mean expression. E) Dotplot of expression of key molecules involved in PLCB2 and PLCG2 upstream activation and downstream signalling in tuft cells and pooled absorptive (TA and enterocytes) and secretory (paneth, goblet and enteroendocrine (EECs)) epithelial cells in the dataset. F) Percent expression of FCγ receptor by SiglecF+EpCAM+ and SiglecF-EpCAM+ cells in wild type mice determined by flow cytometry (n=4). P values were calculated using 2-way ANOVA with Sidak multiple comparisons. ****<0.0001. G) Summary schematic of signalling pathways of tuft cells shown in E.

Focused subclustering of EECs revealed *TPRH1+, NEUROG3-high, NTS/GCG/PYY+, MLN/GHR+, CCK+* and *GAST/GIP+* subsets (Figure 2A & Supplementary Fig. 2B), resembling populations recently described in intestinal organoid experiments ^2^. Comparing the cellular composition of the epithelium across age groups highlighted a drastic developmental shift at approximately 10 PCW, with a majority of undifferentiated epithelial cells prior to this point (Figure 2B & Supplementary Fig. 3B). *NEUROG+* EECs were enriched in the first trimester fetal gut, while *NTS/GCG/PYY+* EECs and *MLN/GHRL+* EECs were only evident from the second trimester through to adulthood (Figure 2B & Supplementary Fig. 3B). *NEUROG3*, a transcription factor that mediates epithelial cell commitment to EEC lineage ^16^, and the absence of mature neuropeptides suggests that the *NEUROG3+* EECs may represent early committed EECs.

M cells, which specialise in transferring luminal antigen through the epithelial lining of the gut were very rare and seen only in pediatric and fetal samples within this dataset (Figure 2B & Supplementary Fig. 3B). Interestingly, expression of MHCII was restricted to postnatal cells (Supplementary Fig. 3C), suggesting that this expression is driven by the presence of the microbiome. All mature epithelial cell types were present from 12 PCW, except paneth cells, which appeared postnatally (Figure 2B & Supplementary Fig. 3B).

The gut epithelium represents a point of viral entry, and SARS-CoV-2 has been shown to enter enterocytes via the receptor, *ACE2*, and protease, *TMPRSS2*^17^. We used our developing gut epithelial dataset to investigate whether viral entry is possible at all life stages. Indeed, both ACE2 and TMPRSS2 were expressed by enterocytes in early development, childhood and adulthood, with TMPRSS2 also expressed by BEST4+ enterocytes and M cells (Figure 2C & Supplementary Fig. 3D). This highlights for the first time the ability for SARS-CoV-2 to infect the fetal gut epithelium.

IBD pathogenesis is due to impaired intestinal epithelial cell function (as well as exacerbated immune functions). We sought to investigate how IBD risk may be linked to cell types across the anatomical regions of the gut, by analysing expression of IBD-associated epithelial cell type marker genes. We focused on the pediatric and adult epithelial compartments, since IBD onset is most prevalent at later life stages. Expression of IBD-GWAS genes was largely consistent across regions, with most variability arising from the presence or absence of cells per region (Figure 2D). In particular, paneth cells present only in the small intestine during health and well documented as metaplastic in colon of IBD patients^18^, showed specific expression of intelectin, *ITLN2*, also previously shown to be enriched in the small intestine ^19^. Surprisingly, *PLCG2*, two missense variants of which have been linked to aberrant B cell responses in early onset IBD ^20,21^ and primary immune deficiency ^22^, was specifically expressed by tuft cells.

To more deeply explore the relevance of *PLCG2* expression in epithelial cells of IBD patients, we compared epithelial *PLCG2* expression across different developmental stages and Crohn’s disease. We observed a small increase in expression of *PLCG2* in Crohn’s disease epithelium (Supplementary Fig. 3E). To further confirm the role of epithelial *PLCG2* in inflammation, we stimulated primary tissue derived intestinal organoids with inflammatory cytokines TNF-alpha or IFN-gamma and performed scRNAseq. While there was no morphological difference in stimulated organoid lines (Supplementary Fig. 3F), we confirm an increase in *PLCG2* expression across stimulated organoid epithelial cells (Supplementary Fig. 3G, n=3).

The discovery of expression of *PLCG2* by tuft cells at higher levels than B and myeloid cell lineages (Supplementary Fig. 3H-J) led us to wonder whether tuft cells specifically could be responsive to signals from immune cells. We therefore analysed the expression of genes involved in immune surveillance by these cells. We screened for expression of ITAM-linked receptors that may activate PLCG2 (Supplementary Fig. 3K). Among these, only *FCGR2A*, which is activated in response to IgG and shown to be expressed by epithelial cells in immunized mice ^23^, was specifically expressed by a fraction (2.75%) of tuft cells (Figure 2E). Using the mouse as a model, we confirmed expression of Fcgr3 (ortholog of FCGR2A) at protein level by approximately 5% of small intestinal tuft cells (Figure 2F & Supplementary Fig. 3L). Receptor tyrosine kinases were also expressed across tuft cells and other epithelial cell types, however since these are known to be mainly linked to PLCG1 activation, whether they are also responsible for PLCG2 activation in tuft cells is difficult to delineate (Figure 2E,G & Supplementary Fig. 3J).

Tuft cells are known to be receptive to chemical signals with essential roles in pathogen responses of the gut ^24–27^. Indeed we confirm expression of the succinate receptor, *SUCNR1*, and downstream mediators *GNAT3, PLCB2, ITPR2, PRKCA* and *TRPM5* (Figure 2E,G), which lead to IL-25-mediated type 2 immune responses. While mouse studies have also implicated TAS1R3 ^28^, we additionally show expression of *TAS1R1*, albeit at a lower level. TAS1R1 forms a heterodimer with TAS1R3 for umami sensing to activate the same type 2 immune signaling ^26^ (Figure 2E,G).

Our in depth analysis of the gut epithelium adds resolution to its changing composition throughout life, showing regional patterning of IBD-GWAS genes in the adult gut and discovering the likely capacity of tuft cells to sense IgG through PLCG2 activation. This leads us to hypothesize that PLCG2 genetic variants may block the receptiveness of tuft cells and inhibit their downstream signaling to other immune cells, thus contributing to increased risk of IBD.

### Enteric nervous system differentiation during development

Similar to the epithelium, the enteric nervous system displayed a shift in development at 10-12 PCW. This provided us the opportunity to investigate the differentiation of human Enteric Neural Crest Cell (ENCC) progenitors. ENCC progenitors are derived from vagal or sacral neural crest cells that colonise the gut during embryonic development. ENCCs must balance proliferation and differentiation into glia and neurons without depleting the precursor pool, as premature differentiation of ENCCs leads to aganglionic colon ^29,30^.

Cycling ENCC progenitor cells, co-expressing *RET, SOX10, EDNRB*, were most evident in first-trimester development (Supplementary Fig. 4A, B). The remaining cycling populations (G2M stages) showed polarised expression of glial and neuronal genes (Figure 3A-B; Methods), suggestive of distinct proliferating committed progenitors. Both of these committed progenitor populations expressed genes distinct from those observed in differentiated cells of ENCC-lineage. For example, committed glial progenitors expressed high levels of *ASPM* and *SGO2* (Supplementary Fig. 4C). Amongst genes expressed by neuronal committed progenitors were *GDF10*, which acts through TGFβ signalling to activate axon sprouting ^31^, multiple Notch ligands (*DLL1, DDL3*) and Notch enhancer *MFNG*, altogether implicating NOTCH and TGFβ signalling in human neuroblast differentiation (Supplementary Fig. 4C). The developmental transition from ENCC progenitor to glial or neuronal progenitors towards terminal cell types was supported by trajectory analysis (Figure 3C, Supplementary Fig. 4D-F).

**Figure 3:**
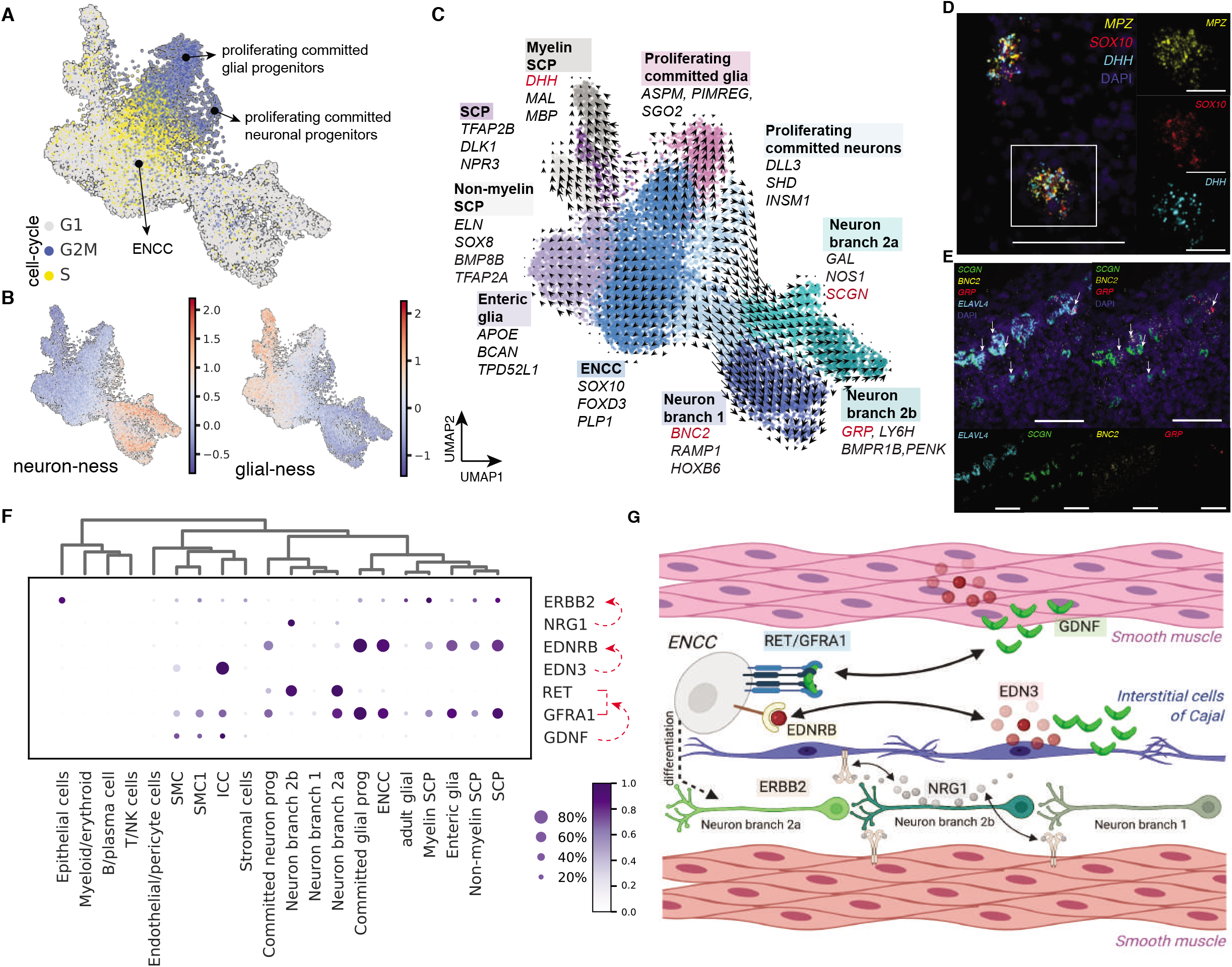
Hirschsprung’s disease risk gene expression in developing human enteric nervous system. A) UMAP projection of fetal enteric neural crest cell progenitors and their progeny coloured by cell cycle score. B) UMAP projection as in A) colored by the glial (left) or neuronal (right) score. Scores were calculated using glial (*SOX10, MPZ, S100B, ERBB3, PLP1, GAS7, COL18A1, EDNRB*) and neuronal (*ELAVL3, ELAVL4, TUBB2B, PHOX2B, RET, CHRNA3*) genes. C) UMAP projection as in A) colored by cell type. The overlaid arrows show scVelo derived differentiation trajectory. Selected marker genes are listed for each population. Multiplex smFISH visualisation of D) myelin Schwann-like cells (scale bars: 100 μm main panel, 30 μm zoom panels), E) three neuronal subtypes described in the fetal 15PCW ileum (scale bars main and zoom panels 100 μm). Arrows highlight three subsets of *ELAVL4+* cells expressing either *SCGN* (greeen), *GRP* (red) or *BCN2* (yellow) genes. F) Dotplot with key Hirschsprung’s disease-associated ligand-receptor genes in the whole fetal dataset. TF= transcription factor, Prog = progenitor. G) Summary schematic of signalling involved in Hirschsprung’s disease as in F.

While ENCCs were abundant in the first-trimester, differentiated neuronal and glial cell types were enriched in the second-trimester of development (Supplementary Fig. 4B). All terminal glial cells expressed high levels of neural crest precursor genes including *FOXD3, MPZ, CDH19, PLP1, SOX10, S100B* and *ERBB3*, but lacked *RET* (Supplementary Fig. 4C). We further defined enteric glia (marked by *BCAN, APOE, CALCA, HES5, FRZB*), and three subsets of Schwann cell precursor (SCPs) all marked by *PLP1, PMP22, CDH19, ITGA4 and CDH2*, but varying in expression of *TFAP2A, DHH* and *FABP7* (BFABP) (Figure 4C). All three subsets lacked expression of *POU3F1* (OCT6), but highly expressed *CDH19*, a marker restricted to SCPs ^32^, supporting their identity as precursors rather than mature Schwann cells. In particular, the first subset, which we labelled ‘SCP’, was enriched in the first-trimester and marked by specific expression of *TFAP2B* and *CXCL13*. Both transcription factor *TFAP2A* (AP-2α) and its paralog *TFAP2B* (AP-2β) have been shown to be expressed in SCPs, but not immature Schwann cells, and regulate timing of SCP differentiation ^33^. The second subset, termed ‘Myelin Schwann-like’, highly expressed myelin associated genes (*MGP, MBP, DHH, NTRK2, SBSPON*), while the third subset, termed ‘Non-myelin Schwann-like’, was enriched in *TFAP2A, ELN, FABP7 and BMP8B* expression (Figure 3C, Supplementary Fig. 3C). We visualised *BMP8B+* Non-myelin Schwann-like cells in the myenteric plexus (Supplementary Fig. 4G), while *DHH+* Myelin Schwann-like cells were found primarily in the mesentery (Figure 3D). SCPs give rise to 20% of enteric neurons with SCP-derived neurons emerging postnatally in mice ^34^. Consistent with this observation, we did not observe differentiation of SCPs into neuronal subsets during embryonic and fetal development in humans (Figure 3C, Supplementary Fig. 4D-F).

**Figure 4:**
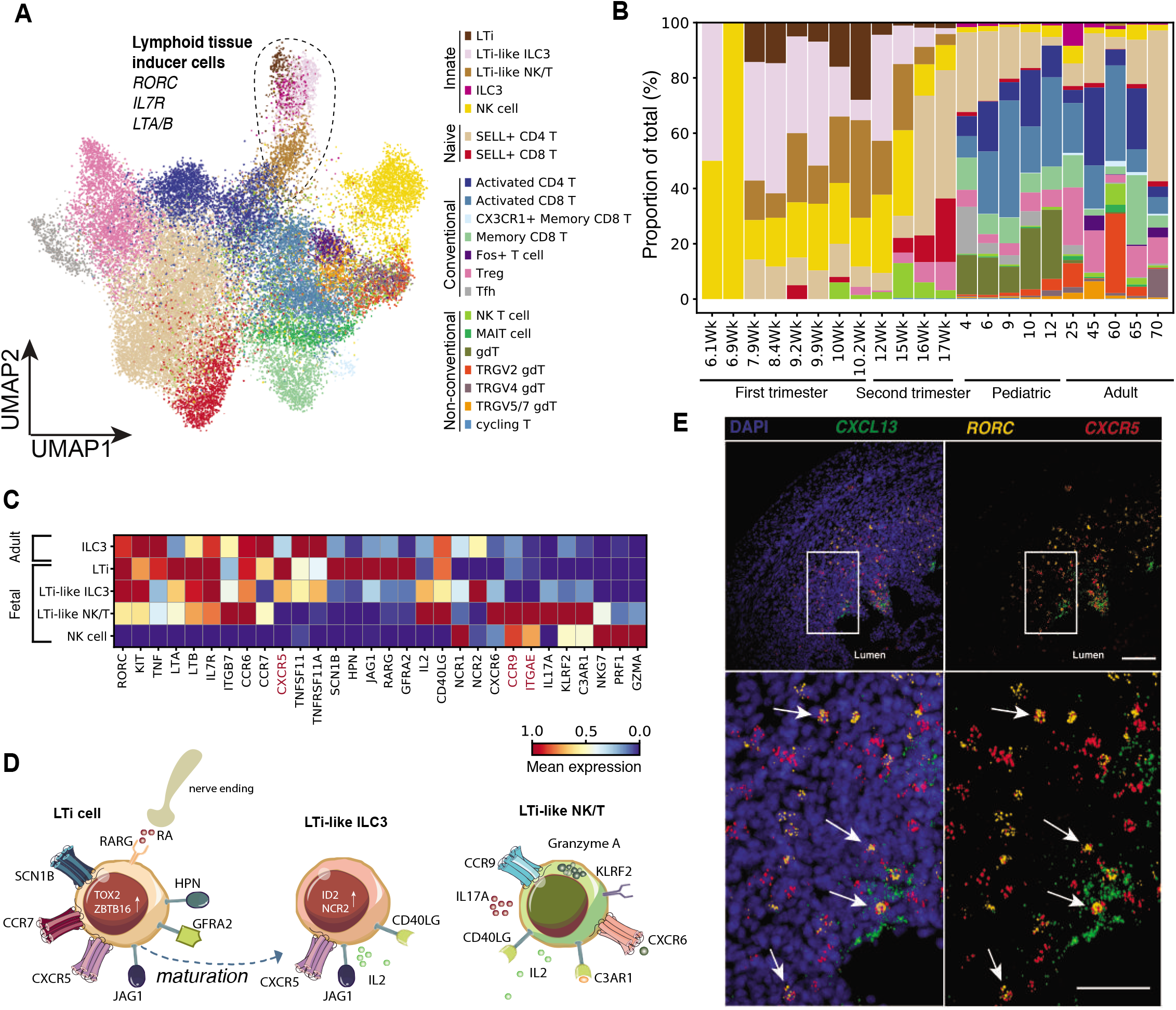
Lymphoid tissue inducer cells in gut-associated lymphoid tissues. A) UMAP visualization of T and innate-like cells in fetal, pediatric and adult samples. Dotted line denotes lymphoid tissue inducer (LTi)-like cell types. B) Bar graph showing the relative proportion (%) of cell types among total T and innate-like lymphocyte population as in A across developmental timepoints. C) Relative expression of key defining and novel marker genes expressed in lymphoid tissue inducer cell types as compared to NK cells. D) Schematic of three LTi-like populations and their features described in this study. E) Multiplex smFISH staining of *CXCL13* (green), *RORC* (yellow) and *CXCR5* (red) in fetal ileum tissue at 15 PCW. The boxed area is shown at higher magnification below. Arrows highlight *CXCR5+RORC+* co-expressing cells. Scale bar main panel=100 μm, zoom panel=50 μm.

Neuronal differentiation extended from neuroblasts into two main neuronal branches, and one additional sub-branch (Figure 3C). Both main branches were found in all intestinal regions (Supplementary Fig. 4H). The first branching event divided branch 1, expressing *BNC2* and *RAMP1*, from branch 2, expressing *ETV1* and *KCNJ5*. Branch 2 further divided into sub-branches 2a and 2b, distinct in expression of gastrin releasing peptide *GRP*, transcription factor *NEUROD6* in sub-branch 2a versus calcium binding protein *SCGN, n*eurexophilin-2 *NXPH2* and netrin-G1 *NTNG1* in sub-branch 2b (Supplementary Fig. 4C). We visualise and validate the cells belonging to branch 1, 2a and 2b in the human myenteric plexus using smFISH in Figure 3E.

We sought to relate developing neuron branches to mature enteric neuron subtypes (Supplementary Fig. 4J). Branch 1 neurons, in addition to *BCN2*, expressed *TSHZ2, ALK* and *HTR4* reminiscent of putative excitatory motor neurons ^7^. Branch 2a neurons expressed genes for both putative inhibitory motor neurons (*NOS1, GFRA1, PTGIR, KCNJ5, DGKB*, and *NPY*) and putative secretomotor/vasodilator neuron markers (*VIP, ETV1, SCGN*, and *UNC5B*) ^7^, suggesting this branch gives rise to both types of neurons. Branch 2b highly expressed *TAC1, PENK, BMPR1B, ADRA2A and NXPH2* reminiscent of putative interneurons ^7^. The developing branches expressed low levels of putative sensory neuron marker genes (*CALCA, NMU, SST, DLX3*), suggesting that this subtype develops later in gestation (Supplementary Fig. 4J).

### Cellular circuitry of Hirschsprung’s disease

Hirschsprung’s disease is characterised by loss or impaired differentiation of the enteric nervous system and can manifest at variable locations of the distal bowel. To identify cell types involved in Hirschsprung’s disease, we screened ENCC-lineage cells for the expression of Hirschsprung’s disease-associated genes ^35,36^. We further included genes associated with Hirschsprung’s disease phenotype in mice^37^, but currently lacking clinical association in humans.

The majority of Hirschsprung’s disease-associated genes were expressed across multiple differentiating populations with varying intensity (Supplementary Fig. 4I). For example, *UBE2T* was most highly expressed by committed proliferating progenitors, while other Hirschsprung’s disease-associated genes were highly expressed in glial (*ZEB2, COMT, NTRK3*) or neuronal (*TLX2, SEMA3D, L1CAM, HOXB5*) compartments (Supplementary Fig. 4I). Expression levels of disease genes also varied between the three developing neuron clusters (branches 1, 2a and 2b). For example, *ASCL1* was highly expressed in branch 1 and 2a, but not in branch 2b. *RET* was selectively absent from branch 1, and *GFRA2* absent from branch 2a.

Interestingly, *CDH2, L1CAM* and *RET* were more highly expressed in colonic neuronal lineage cells compared to the equivalent cells in the small intestine, while the opposite was true for *ATRX* and *ASCL1* (Supplementary Fig. 4I). We also observe unexpected expression of *NTRK3* in pro-myelin SCPs (Supplementary Fig. 4I). Mice lacking Ntrk3 or its ligand Nt3 have reduced numbers of intrinsic sensory neurons ^38^, suggesting that pro-myelin SCPs may support the differentiation of neurons during development.

To understand the contribution of mesenchymal cell subsets to pathogenesis of Hirschsprung’s disease, we investigated the expression of key Hirschsprung’s disease-associated ligands and their receptors ^35,36^ across gut cells of all developmental ages (Figure 3F). Among these ligands was *GDNF* that activates *RET* and its co-receptor *GFRA1, EDN3* and its receptor *EDNRB3*, and *NRG1 a*nd its receptor *ERBB2*. We observed expression of *GDNF* and *EDN3* primarily by smooth muscle cells (SMC) and interstitial cells of Cajal (ICC) (described among stromal cells below; Figure 3F). While in adult gut *NRG1* is expressed by glial and multiple neuron subtypes ^39^, in the fetal gut *NRG1* was highly expressed in neuronal branch 2b. Its receptor *ERBB2* was expressed across glial and stromal cells including ICC and SMC (Figure 3F,G). *NRG1* acts as a neurotrophic factor and supports growth, proliferation and differentiation of the enteric nervous system. Our analysis implicates fetal branch 2b neurons, and ICC/SMC interactions, in the pathogenesis of Hirschsprung’s disease.

### Formation of secondary lymphoid structures

The gut has its own elaborate nervous system as detailed above, and is also a unique organ with respect to having its own immune system. Secondary lymphoid structures (SLO) are key sites of immune surveillance and serve to initiate adaptive immune response to exogenous pathogens. Development of SLOs in the gut, including gut-associated lymphoid tissues (GALT) and mesenteric lymph nodes (mLN), has been reported to begin between 8 to 12 PCW in humans ^40^. We observed mLNs starting around 12 PCW, with structures clearly recognisable and disectable at 15 PCW (Supplementary Fig. 5A). In order to better understand the fetal gut-resident immune cells and their organisation in humans, we investigate in detail the key cell types involved in lymphoid tissue organisation during embryonic development.

Lymphoid tissue inducer (LTi) cells, categorised as group 3 innate lymphoid cells (ILC3), interact with lymphoid tissue organiser (LTo) cells to initiate SLO formation during development in mice and humans^41^. Sub-clustering of fetal and adult T and innate lymphoid cells revealed three cell clusters matching published characteristics of LTi cells (Figure 4A,B). These characteristics include high expression of *RORC, KIT, NRP1, TNF, LTA/B, IL7R*, and *ITGB7* (integrin α4β7) (Figure 4A,C) as well as the absence of productive αβ TCR chains (Supplementary Fig. 5B). The cluster labelled innate lymphoid cell progenitor (ILCP) was further defined by high expression of chemokine receptors *CXCR5* and *CCR7*, and high expression of cell adhesion molecule *SCN1B* ^42^ and serine protease encoded by *HPN* gene (Figure 4C). The second cluster labelled ‘LTi-like ILC3’ clustered closely with adult ILC3 cells and had the highest expression of *TNFRSF11A* (RANK) and its ligand *TNFSF11* (RANKL), as well as *NCR1* (NKp46) and *NCR2* (NKp44) supporting an ILC3 phenotype for this cell population (Figure 4C). Both ILCP and LTi-like ILC3 cells expressed high levels of *NRP1*, previously shown to mark human ILC3 cells with LTi cell function ^43^.

Lastly, a third population, which we term ‘LTi-like NK/T’, express *IL17A, ITGAE* and *CCR9*, and NK-associated genes including *NKG7, PRF1, and GZMA*, but lack expression of *CXCR5*. While we did not capture productive TCRαβ chains in this population, we observed expression of CD3 genes (*CD3G, CD3D and CD3E*) and TCRδ genes (*TRDC, TRDV2*), suggesting they are γδ T cells (Figure 4C, Supplementary Fig. 5B). Reassuringly, these three LTi-like cell subtypes were also identifiable in our full-length scRNAseq data from fetal ileum cells (Supplementary Fig. 5D).

ILCP cells were enriched at 6-10 PCW, while LTi-like ILC3 and LTi-like NK/T cells were present at 6-17 PCW (Figure 4C) and expanded in all gut regions throughout development (Supplementary Fig. 5E). Only ILCP cells were captured in fetal mLN (Supplementary Fig. 5E), suggesting that LTi-like subsets are expanded during GALT, but not mLN development. This observation together with expression of key chemokine signals (*CXCR5, CCR7, CCR6*) and RNA velocity analysis suggested that ILCP cells are the first LTi-like cells in the developing gut. Hence they likely represent a progenitor state to mature ILC3s (Figure 4C, Supplementary Fig. 5F). RNA smFISH staining placed *CXCR5+ RORC+* ILCP/LTi-like ILC3 cells in proximity to *CXCL13+* LTo cells in fetal proximal gut mucosa, supporting the concept of congregation of these cells beneath the epithelium in the developing gut (Figure 4D).

In order to infer whether the developing lymphoid structures were partitioned into B and T cells zones as evident in mature lymphoid tissue, we characterised B cells in fetal and adult tissues. During fetal development, common lymphoid precursor (CLP) cells (*CD34, FLT3, SPINK2*), Pro-B (*DNTT, VPREB1/3, IGLL1, RAG1/2*), Pre-B (*CD19, CD38, IGLL5, RAG1/2*) and immature B cells (*CD40*, productive IgM, MHC II) were evident (Supplementary Fig. 6A-E). In the adult gut, we detected light zone (*EBI3, LMO2, GMDS* and *BCL2A1*) and dark zone (*AICDA, SUGCT, EZR* and *ISG20*) germinal centre B cells enriched in the pediatric terminal ileum. This supports the presence of fully mature and organised lymphoid aggregates in the adult but not fetal tissue (Supplementary Fig. 2B, 6F-G).

Analysis of paired V(D)J sequencing data revealed almost exclusive IgM heavy chain expression across all fetal gut B cells and no evidence for meaningful levels of clonal expansion and somatic mutation. In comparison, adult B cells expressed primarily IgA1 and IgA2 with minor populations of IgM, IgG1 and IgG2 cells, fitting with our previous report ^6^, and displayed high mutational frequency and clonal expansion consistent with having undergone affinity maturation (Supplementary Fig. 6H-I). Mutation frequency and clonal expansion were slightly higher at the proximal and distal ends of the adult gut, and sharing of B cell clones, while restricted to occurring within each donor, was evident across distant gut regions (Supplementary Fig. 6J-L). These results suggest zonation of B cells within postnatal gut mucosa, in addition to antigen activation, class-switching of B cells and dissemination of B cell clones between small and large intestine.

In contrast to adult B cells, we found no evidence of fetal B cell expansion or maturation in response to antigen. This suggests that additional environmental factors, such as microbiota, might be required for the maturation and zonation of B cells in secondary lymphoid organs. Taken together, our data identifies three types of LTi cells orchestrating the earliest development of lymphoid structures, and B cells that are poised for response in these sites in the prenatal intestine.

### Stromal cells in human mLN and GALT development

LTo cells are LTβR+ non-hematopoietic cells that interact with LTi cells during lymph node formation, and can be either mesenchymal (mLTo) or endothelial (eLTo) in origin. LTi and LTo cell interaction leads to activation of canonical and non-canonical NF-κB pathways, and results in LTo secretion of chemokines CCL19, CCL21 and CXCL13 as well as expression of adhesion molecules VCAM-1, ICAM-1 and MAdCAM-1. After visualising *RORC+ CXCR5+* LTi-like ILCP/ILC3 cells in proximity to *CXCL13+* cells in fetal gut mucosa as mentioned above (Figure 4D), we postulate that GALT formation follows similar mechanisms to those previously described in the mLN. To address this, we analyse stromal populations from both gut tissue and mLN.

Within the entire dataset, we identify multiple stromal populations defined by the expression of *VIM, COL1A2* and *COL6A1* (Supplementary Fig. 2B) including myofibroblasts, smooth muscle cells (SMC, SMC1), pericytes, ICCs, mesothelium, as well as populations resembling stromal cells previously described in the colon (labelled as S1-S4) ^8^ (Figure 5A). We further identified fibroblast populations typically defined in mouse lymph nodes ^44^, including T reticular cells (expressing *CCL21, CCL19* and *GREM1*) and follicular dendritic cells (FDCs; expressing *CXCL13, CR1* and *CR2*) (Supplementary Fig. 7A). T reticular cells and FDCs were found in the terminal ileum (Supplementary Fig. 7B). These samples were also enriched for LZ and DZ GC B cells (Supplementary Fig. 2B, 6F-G), suggesting the presence of cells that organise Peyer’s patches.

**Figure 5:**
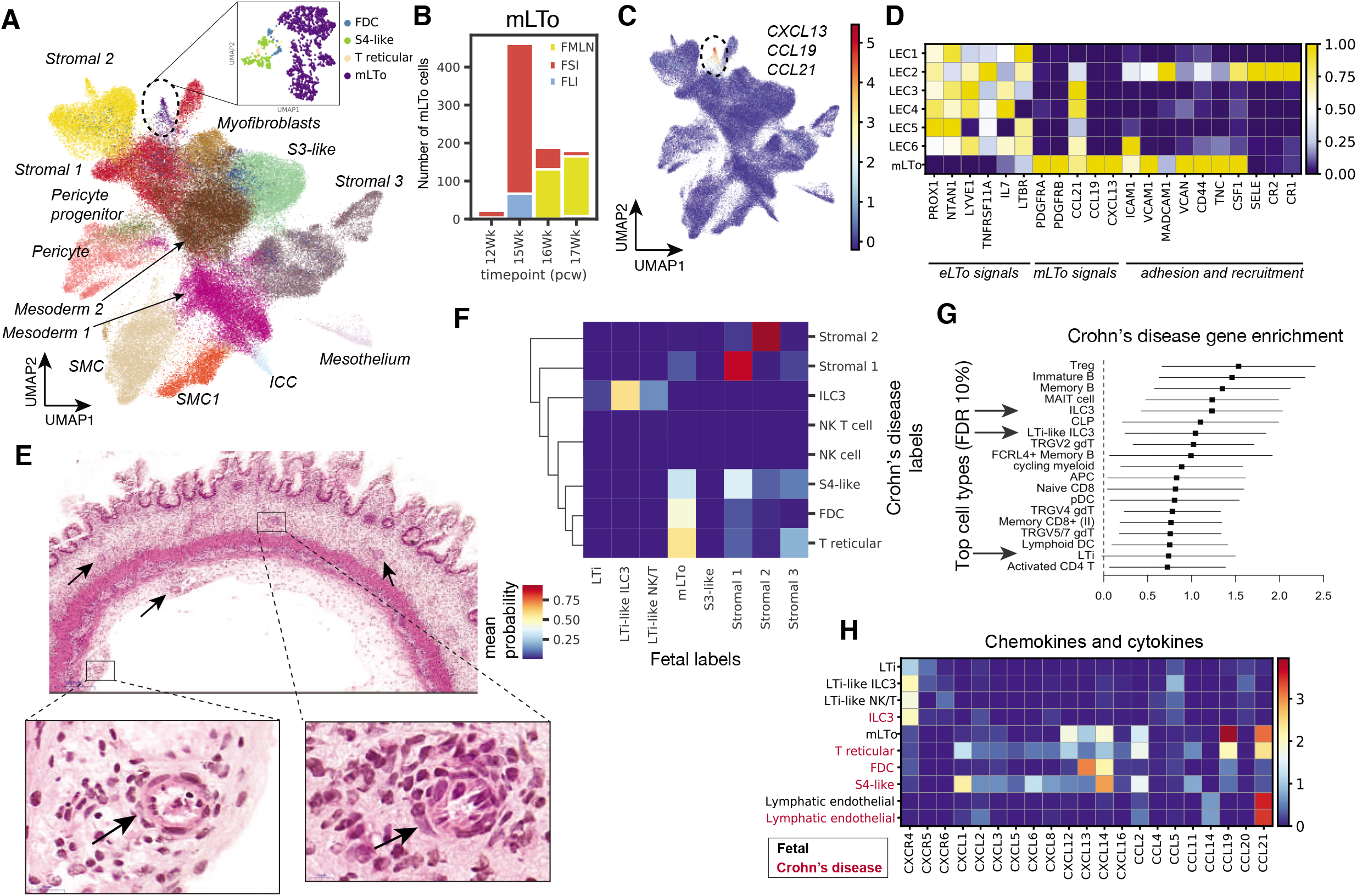
Lymphoid tissue organogenesis programs adopted in Crohn’s disease. A) UMAP projection of fetal, pediatric and adult stromal cells colored by cell type. Zoom in panel shows mLTo cells, S4-like, follicular dendritic (FDC), and T reticular cells. B) Bar plot showing number of mesenchymal lymphoid tissue organiser cells (mLTo) captured in the dataset and colored by gut region. C) Feature plot with cells coexpressing *CXCL13, CCL19*, and *CCL21*. D) Heatmap showing scaled expression of selected genes involved in lymphoid tissue organisation in model organisms. E) H&E staining of cross-section of fetal colon (15 PCW). Magnified panels show developing vessels. F) Heatmap showing mean probability of cell types matching between fetal and Crohn’s disease stromal cell populations. G) Top cell types across fetal, pediatric, and adult data enriched in genes associated with Crohn’s disease (FDR 10%). H) Heatmap with expression of cytokines and chemokines in cells involved in tertiary lymphoid organ development of fetal (black) and functionally related cell types in pediatric Crohn’s disease (red).

In prenatal intestine (12-17 PCW) and mLN, we observe a stromal cell population transcriptionally similar to T reticular, FDCs and S4 fibroblasts (Figure 5A subpanel, 5B & Supp. Figure 7C). This stromal cell cluster, absent from pediatric and adult samples, was marked by expression of chemokines *CCL19, CCL21* and *CXCL13* as well as adhesion molecules (Figure 5C), and resembles activated mLTo cells described in developing mouse lymph nodes ^45^. Other genes expressed in mLTo cells were components of the NF-κB pathway (*NFKBIA, NFKB1/2, IL32, REL, RELA/B*), S1P transporter spinster homolog 2 *SPNS2*, involved in T and B cell trafficking ^46^ and chemokines *CXCL2* and *CXCL3* (Supplementary Fig. 7D).

We further investigate the transcriptional differences of mLTo cells from lymph nodes and intestinal regions to define differences that could lead to differential positioning of SLOs. Interestingly, large intestinal mLTo cells share transcriptional characteristics with mLN mLTo cells, including expression of *C7* and *SLC22A3* (Supplementary Fig. 7E), suggesting similarities between mLN and cryptopatch development. Amongst the genes highly expressed in mLN mLTo cells were angiotensin receptor *APLNR*, endothelin receptor *EDNRA* and extracellular matrix protein *TNC*, all with roles in blood vessel formation. In addition, mLTo cells in mLNs expressed *IL33* and *FDCSP*, also present in lymph node and spleen fibroblastic reticular cells ^47^ and observed in fibroblasts expanded in ulcerative colitis ^8^. Thus, transcriptionally similar mLTo cells are implicated in the formation of human GALT as well as mLN tissue development.

### Endothelial cells in mLN and GALT organisation

As lymphatic endothelial cells (LECs) connect the mLN to the tissue and can contribute to SLO formation by interacting with LTi and mLTo cells, we investigated endothelial cell heterogeneity. Amongst endothelial cells we observe arterial, venous, capillary and lymphatic endothelial cells (Supplementary Fig. 2B, 7F-G). LECs separated into six clusters (labelled LEC1-6), all uniformly expressing *PROX1* and *LYVE1*, and *TNFRSF11A* (RANK), *IL7* and *CCL21-* signals required for initial recruitment of LTi cells (Figure 5D; Supplementary Fig. 7H,I). LEC1 and LEC5 ressemble subsets described in human lymph nodes ^48^. The LEC1 subset is marked by specific expression of *C7*, *TMEM100* and abundant expression of *ACKR4* and *CAV1*, resembling cells lining the subcapsular sinus ceiling in the human lymph nodes ^48^, while LEC5 express *CLDN11, ANGPT2* and *GJA4*, shown to be expressed on collecting lymphatic valves (Supp. Figure 7I)^48^. We further define LEC3 present mostly in the pediatric and adult intestinal regions, and LEC4 specific to developing intestinal regions (Supplementary Fig. 7J), which may represent developing and differentiated lymphatic vessels present in the intestinal mucosa.

LEC2 cells emerge during the second trimester (15-17 PCW) and are present in gut regions as well as fetal mLNs (Supp. Figure 7J). This population expresses *TNFRSF9, THY1, CXCL5*, and *CCL20*, similarly to a population of LECs lining the subcapsular sinus floor in human lymph nodes (Supp. Figure 7I)^48^. In addition, they express genes *ANO9, PSD3* as well as high levels of adhesion molecules including *MADCAM1, VCAM1* and *SELE*, suggesting their involvement in lymphocyte recruitment (Figure 5D, Supplementary Fig. 7K). LEC6 was marked by expression of *ADAMTS4*, metallothionein genes such as *MT1X, MT1E*, and is enriched in the adult appendix and mLN (Supplementary Fig. 7I,J). Similarly to mLTo cells, LEC2 and LEC6 express targets of the canonical NFkB pathway (Supplementary Fig. 7L). As LEC2 is present during development, NFkB pathway activation suggests that the LEC2 population contributes to organisation of GALTs. Finally, histological staining of fetal ileum allows us to identify blood vessels resembling high endothelial venules (Figure 5E), which would permit entry of lymphocytes into the tissue.

Mesenchymal and endothelial cells are central to the recruitment and positioning of leukocytes in developing and mature mLN and GALT structures. To determine cell-cell interactions governing early lymphoid structure formation and leukocyte recruiting, we investigate complementary ligand-receptor expression across LECs, mLTo and LTi-like subsets. Amongst ligand-receptors expressed were *LTA* and *LTB* (LTα1β2) expressed on all LTi-like cell subsets and *LTBR* (LTβR) on mLTo and multiple LEC subsets (Supplementary Fig. 8A), characteristic of LTo and LTi cells ^41^. Interestingly, we found that LTi-like ILC3 and NK/T cells utilise CD40LG to signal to mLTo and LEC through either *CD40* or integrin a5b1 (*ITGA5*), respectively (Supplementary Fig. 8A). The activation of this signalling pathway in endothelial cells has been previously reported to drive angiogenesis ^49^, suggesting that LTi-LEC interaction may further enhance local vascularisation. *IL2* was expressed by LTi-like ILC3 and NK/T cells and its receptor *NFGR* was expressed specifically on mLTo cells. This ligand-receptor interaction may facilitate LTi support for mLTo proliferation and survival in GALTs (Supplementary Fig. 8A). In addition, cognate receptor expression was evident across immune cell types that would enable their recruitment into the developing SLO (Supplementary Fig. 8B). For example, mLTo cells also expressed *CCL2* for recruitment of monocytes and dendritic cells *via* their receptor *CCR2* (Supplementary Fig. 8B).

### SLO developmental programmes are adapted during Crohn’s disease

SLO development is pre-programmed and does not require antigen stimulation. In contrast, ectopic lymphoid structures, organised aggregates of lymphoid cells, develop in response to chronic inflammation ^50^. These ectopic structures often involve T and B cell segregation and increased vascularisation. Compared to SLOs, ectopic lymphoid structures are often transient and resolve after antigen clearance. Following on from our observations of lymphoid structure formation during development, we compared similarities and differences between normal organogenesis and ectopic lymphoid structure formation often observed in Crohn’s disease (CD)^51^.

We use logistic regression modeling of pediatric CD scRNAseq data ^4^ to calculate the mean prediction probability of matching fetal and pediatric CD cell types (Figure 5F). We observed ILC3 cells in CD that matched fetal LTi-like ILC3 with 60% probability (Figure 5F). S4-like, FDCs and T reticular cells in CD matched fetal mLTo (Figure 5), the former two of which were expanded in four out of seven CD donors (Supplementary Fig. 8C-D). Curiously, an equivalent stromal population has recently been shown to be expanded in ulcerative colitis ^8^. In addition, transcriptionally similar populations are expanded in leukocyte-rich rheumatoid (RA) ^52^ and cirrhotic liver disease^53^ (Supplementary Fig. 8E).

To determine cell types involved in CD, we calculated the enrichment score for CD-associated GWAS gene expression for each cell type and selected top cell types (FDR 10%; Methods). Adult ILC3, and fetal ILCP and LTi-like ILC3 cells were among the top cell types enriched for expression of CD-associated genes (Figure 5G). Finally, we compared cytokine and chemokine expression between fetal and CD counterparts, and showed core genes conserved between these cells (Figure 5H). For example, ILC3 and ILCP/LTi-like ILC3 commonly expressed *CXCR4, CCL5, CCL20*, while *CCL4* was specific to adult ILC3. mLTo and CD fibroblasts showed common expression of *CCL19, CCL21, CXCL12-14* and *CCL2*, while *CXCL1/2/3/5/6/8* and *CCL11* were specific to adult CD cells.

Our identification of equivalent cell types with the same cellular networking in lymphoid aggregate formation in fetal and CD gut suggests a reactivation of the programs for lymphoid formation during disease.

## Discussion

In this study, we present an integrated dataset of over 346,000 single cells of multiple anatomical regions of the human gut throughout development, childhood and adulthood, browsable online at gutcellatlas.org. We discover epithelial and neuronal lineage composition and developmental relationships, and reveal cell-type specific expression of IBD- and Hirschsprung’s disease-associated genes, respectively. We identify the three key cell types of mesenchymal, endothelial and innate lymphoid origins that orchestrate lymphoid organ formation in fetal development and show reinitiation of this cellular program during Crohn’s disease.

In the epithelial compartment, we are able to subdivide enterocytes in unprecedented detail. BEST4+ enterocytes have previously been shown to function in pH sensing and transport of metals, ions and salts, but have been restricted to the colon ^9,14,54^. Here, we show their presence in the small and large intestines, and as early as the first trimester in the small intestines, supporting a consistent requirement for this population along the gut and throughout life. In addition, we screened for expression of the SARS-Cov2 receptor, ACE2, and associated protease, TMPRSS2, in the gut epithelium. We advance on previous observations of their expressed in epithelial cells ^55^ by showing their expression in enterocytes throughout life, including fetal development. Since amniotic fluid and placental tissue of an infected new mother have tested positive for SARS-Cov2 ^56^, this could represent a novel route for viral transmission *in utero*.

Continuing our analysis of the epithelial compartment, we discover that FCGR2A, a receptor activated by the Fc fragment of IgG upstream of PLCG2, and other downstream signaling molecules are expressed by tuft cells. We suggest that this pathway could mediate direct immune sensing by tuft cells. This is surprising since FcG receptors have rarely been observed in non-haematopoietic cells ^57^. To our knowledge, FcgrII expression has only been observed in mouse intestinal epithelial cells after immunization with bacterial neurotoxin ^23^ and by human nasal epithelial cells in response to bacteria-derived ligands ^58^. We also note expression of the FCGR2A by our previously described thymic mTEC(II) epithelial population, which expresses low levels of tuft cell transcription factor POU2F3 (developmentcellatlas.ncl.ac.uk). FcG receptors on immune cells mediate response to local IgG via production of inflammatory signals. Although FCGR2A was only expressed by a subset of tuft cells, we expect this would similarly increase during gut inflammation when luminal IgG levels rise. Overall, this data suggests a novel and potentially very impactful immune-sensing role for intestinal tuft cells.

While epithelial cells are responsible for the major absorptive and secretory functions of the gut, the enteric nervous system is necessary for peristalsis. Absence of the ENS is a very serious condition: Hirschsprung’s disease is a congenital disorder characterised by the absence of nerve cells. To date, the genes associated with developmental conditions such as Hirschsprung’s disease have been mapped in humans using adult cells ^59^. We leverage the fetal cells of this dataset to map the expression of Hirschsprung’s disease-associated GWAS genes, and show expression of a set of genes including *RET* and *EDNRB* in multiple enteric neuronal cell types in early human gut development. We also discover for the first time a signaling circuitry amongst ENS and ICC and muscle cells involving Hirschsprung’s disease genes. These observations help explain the difficulty linking HSRC-associated genes with neuronal cell types in mature gut tissue.

Like the ENS, the immune system of the gut is both sophisticated and essential to proper functioning and maintenance of this barrier tissue. Recent single-cell studies have elucidated the diversity and activation of immune cells in the developing human gut ^60,61^. Significant species differences exist in location and size of the SLOs ^62^, major hubs of immune cell surveillance and activation. However, our understanding of the formation of these structures is mostly derived from animal models. As such, using genetically perturbed mice, a three-cell model was proposed for SLO formation ^41^. Here, we identify cell types with characteristics of the three populations: LTi, mLTo, and eLTo (LEC) cells in the developing mucosa and mLN, suggesting a similar process occurs in humans. We provide detailed characterisation of subtypes of the LTi-like cells in both fetal and adult tissues, describing three different subsets of these cells. Based on previous *in vitro* studies, where *RORC*-expressing CD56+ CD127+ IL17A+ cells ^63^ and ILC3 cells ^43^ have been both shown to activate mesenchymal cells, we propose that all three of the LTi-like subsets defined in this study act in lymphoid tissue initiation.

Previous studies point to immune cell programs that support normal development, however, when overactivated, can lead to inflammation in early life such as necrotising enterocolitis and IBD^61^. Here we describe SLO formation as one such program adopted in chronic inflammation to recruit and retain immune cells at the site of active inflammation. Using direct comparison of LTi subsets, mLTo and eLTo cells in development and CD, we identify conserved chemokine and cytokine signals between these populations, as well as point to CD specific signals expressed by ILC3, stromal and endothelial cells in disease.

Expansion of an inflammatory fibroblast population, termed Stromal 4 cells, has been reported in ulcerative colitis patients ^8^ and activated fibroblasts have been reported in adult and pediatric CD ^4,13^. Interestingly, we further observe two specialised fibroblast populations in pediatric CD, namely FDCs and T reticular cells. Similar cell types have been described in mouse lymph nodes ^44^, but have not been captured in the gut mucosa. Together with the presence of light zone and dark zone B cells in the gut (Supplementary Fig. 2B, 6F-G), this suggests the presence of stromal populations involved in SLO zonation as observed in Peyer’s patches previously ^64–67^.

Direct comparison of these cells in development *versus* disease allowed us to separate out expression programs driving SLO formation in development or disease. We show that mLTo-like fibroblasts in CD have unique expression of cytokines and chemokines (e.g *CXCL2-8*) associated with increased recruitment of monocytes and neutrophils to the site of inflammation that may be driving ectopic lymphoid formation. In fetal LTi-like cells, we observed expression of *IL2* and *IL17A*. These inflammatory cytokines may also contribute to controlled inflammatory reaction in the developing fetus, aiding establishment of organised lymphoid tissues prior to birth ^68^. While others have proposed that small numbers of LTi-like NK cells are present in post-natal tonsils ^63^, we do not observe ILCPs or LTi-like NK/T cells in the pediatric or adult gut. Rather, we show transcriptional similarities between ILC3 and fetal LTi-like ILC3 cells. It is possible that ILC3 in CD may act as an initiator cell to activate the mesenchymal cells and recruit other lymphocytes to the site of inflammation.

Overall, our detailed cell annotation of the human intestine across time and space can be mined for cell-type expression of genes associated to any number of intestinal tract diseases. For instance, our identification of cellular programs of prenatal SLO development reactivated during ectopic lymphoid tissue in CD may represent a target for controlling inflammation during this disease, and could also be extended to other inflammatory diseases characterised by ectopic lymphoid structure formation ^69^. Finally, this work opens avenues for using principles of prenatal development and differentiation trajectories outlined here for stromal, epithelial, ENS and immune compartments, to engineer human cells and guide cell maturation *in vitro*, for the purposes of basic research or even screening, and ultimately as therapeutics for regenerative medicine.

## Acknowledgements

We acknowledge the support received from the Wellcome Sanger Cytometry Core Facility, Cellular Genetics Informatics team, Cellular Generation and Phenotyping (CGaP) and Core DNA Pipelines. This work was financially supported by the Wellcome Trust (WT206194, S.A.T.); the European Research Council (646794, ThDefine, S.A.T.); an MRC New Investigator Research Grant (MR/T001917/1, M.Z.); and a project grant from the Great Ormond Street Hospital Children’s Charity, Sparks (V4519, M.Z.). The human embryonic and fetal material was provided by the Joint MRC / Wellcome (MR/R006237/1) Human Developmental Biology Resource (www.hdbr.org [hdbr.org]). We thank Professor Roger Barker, Xiaoling He, Alexander Ross and Steve Lisgo for access to and handling of fetal tissue; David Fitzpatrick for discussion on developmental intestinal disorders; Tong Li and Ola Tarkowska for image processing and infrastructure support; We acknowledge J. Eliasova for the graphical images. We thank the tissue donors and donor families.

## Author contribution

S.A.T, M.H and K.R.J initiated, designed and supervised the project; K.M and K.S.P carried out adult tissue collection; K.R.J, R.E, M.D, S.P, L.B, S.F.V and M.P performed adult tissue processing and scRNAseq experiments; S.L, R.A.B, K.I.G, J.E, C.D supported fetal tissue processing and scRNAseq experiments; L.M, L.B,E.S performed library preparation; A.F performed flow cytometry validation in mice and data interpretation; T.R.W.O, C.E.H, L.S.C provided pathology support; K.R, S.P, and O.A.B performed tissue sectioning, staining and imaging; K.R.J and R.E analysed single-cell data and generated figures; N.K, K.P, S.V.D and N.H provided analysis support and contributed to analysis; H.W.K performed BCR analysis and contributed to data visualisation; E.D performed differential abundance analysis; J.M, M.M, M.K, K.B.M, M.Z, H.U and M.R.C contributed to result interpretation; K.R.J, R.E and S.A.T wrote the manuscript; All authors contributed to discussion and interpretation of results as well as editing of the manuscript.

## Supplementary Figures

**Supplementary Fig. 1: Data quality control.** A) Schematic with tissue processing strategy for second trimester fetal and adult samples. After enzymatic dissociation, either total fraction was loaded onto 10X chip or CD45+/- cell factions were separated using magnetic cell sorting (MACS) and both fractions were loaded on the 10X chip separately. Lymph nodes were processed without enrichment. Second trimester fetal and adult cell samples were processed using 5’ v2 10X kits (Methods). B) Pre-processing and quality control of single-cell RNA-seq data generated in this study and described previously^4^. In short, four datasets, namely first trimester fetal, second trimester fetal, pediatric and adult, were preprocessed separately (including quality control and scrublet doublet removal). Firstly, dimension reduction, clustering and annotation by cell lineage was performed on each dataset separately. Each cell lineage was sub-clustered and fine-grained cell type and state annotation was performed based on marker gene expression. The four datasets were then merged together and each lineage was sub-clustered to unify cell type labels where appropriate. UMAP visualisations show combined dataset colored by sample age, enrichment fraction and donor name.

**Supplementary Fig. 2: Cell types defined in the study and intestinal region variability in BEST4+ enterocytes.** A) Bar plots with relative proportion (%) of cell lineages in each sample grouped by anatomical region within the scRNAseq dataset as in Figure 1B. B) Dot plot for expression of marker genes of cell types and states in each cell lineage in scRNAseq dataset. C) UMAP visualisation of BEST4+ enterocytes colored by key marker *BEST4/OTOP2* expression and region group (fetal and pediatric/adult). D) Volcano plot for differential abundance (DA) between cells from small intestine and large intestine. Each point represents a neighbourhood of BEST4+ enterocytes (FDR: False Discovery Rate, logFC: log-Fold Change). The results for the DA test in adult samples (red) and fetal samples (blue) are shown. The dotted line indicates the significance threshold of 10% FDR. E) Heatmap showing average neighbourhood expression of genes differentially expressed between DA neighbourhoods in Adult BEST4+ enterocytes (1502 genes). Expression values for each gene are scaled between 0 and 1. Neighbourhoods are ranked by log-fold change in abundance between conditions. Genes highlighted in red are specific to small intestinal BEST4+ enterocytes F) Expression of CFTR (antibody: CAB001951) in small intestinal (top) and colonic (bottom) histological sections from Human Protein Atlas (proteinatlas.org).

**Supplementary Fig. 3: Epithelial cell types throughout intestinal life.** A) UMAP projection of detail (left) and pooled pediatric and adult (right) epithelial cells coloured by gut region. B) Bar plots with number of epithelial cells subtypes separated by donor age (row). Units of age is years unless specified as weeks. C) Violin plot with MHCII score in M cells. D) Dot plot of expression of *TMPRSS2* and *ACE2* by epithelial cell types in pediatric (top) and adult (bottom) intestine. E) Bar plot showing mean expression of *PLCG2* in all epithelial cells grouped by developmental/disease phenotype, where each dot is a donor and confidence interval is standard deviation. F) Representative brightfield images of pediatric intestinal organoid line (derived from healthy donors) without (NT) or with stimulation with inflammatory recombinant human proteins IFN-gamma or TNF-alpha. Scale bar = 400 μm. G) Bar plot showing mean expression of *PLCG2* in all intestinal organoid cells grouped by stimulation condition, where each dot is corganoid line (n=3) and confidence interval is standard deviation. H) Heatmap with top differentially expressed genes in tuft cells. The legend indicates whether the gene has known association with tuft cells (purple) or are novel (orange). I) Immunohistochemical staining of HPGDS (antibody: HPA024035), PSTPIP2 (antibody: HPA040944), BMX (antibody: CAB032495), MYO1B (antibody: HPA060144), FYB1 (antibody: CAB025336), SH2D7 (antibody: HPA076728), PLCG2 (antibody: HPA020099) in small intestine from Human Protein Atlas (proteinatlas.org). J) Heatmap of expression of IgG receptors (green) and chemosensing (yellow) and downstream signalling molecules as in Figure 2E for epithelial cells with B and myeloid cells for reference. K) Dotplot showing expression of ITAM- and ITIM-linked receptors and receptor tyrosine kinases across tuft cells and pooled absorptive (TA and enterocytes) and secretory (paneth, goblet and EECs) epithelial cells. L) Representative flow cytometry plots of FcyRII/III staining on EpCAM+SiglecF-(non-tuft epithelial cells) and EpCAM+SiglecF+ (tuft cells) cells and Isotype staining of EpCAM+SiglecF+ cells. Numbers show the percentage of cells within the gate out of the total population.

**Supplementary Fig. 4: Cell types in the developing enteric nervous system.** A) Feature plots with overlaid expression of key genes (*RET, EDNRB, SOX10*) that define enteric neural crest cells (ENCCs). B) Area plot with relative abundance (percentage %) of cell types amongst ENCC-lineage populations as described in Figure 3C across developmental timepoints. C) Dotplot with differentially expressed genes in ENCC-lineage subsets as described in Figure 3C. Genes in red are discussed in text. D) PAGA visualisation of ENCC-lineage cells. Red arrows show ENCC division to committed progenitors. E) UMAP visualisation of ENCC-lineage cells colored by monocle 3 pseudotime. Arrows show differentiation to neuronal branches. F) Heatmap showing genes that change across pseudotime in three neuronal branches shown with arrows in E. G) Multiplex smFISH staining of non-myelin Schwann cells using probes for *BMP8B* (green), *MPZ* (yellow) and *SOX10* (red) in fetal ileum tissue at 15 PCW. The boxed area is shown at higher magnification below. Scale bar main panel=100 μm, zoom panel=30 μm. H) UMAP visualisation of ENCC-lineage cells colored by intestinal anatomical region. I) Heatmap with relative mean expression of genes associated Hirschprung’s disease in humans and ENS defects in mice (not determined). Mean expression was calculated and visualised for small and large intestinal regions separately. J) Dotplot with mature neuron gene expression in developing neuronal branches. PEMN-putative excitatory motor neurons; PIMN-putative inhibitory motor neurons; PSVN-putative secretomotor/vasodilator neurons; PIN-putative interneurons; PSN-putative sensory neuron

**Supplementary Fig. 5: Identification of LTi cell-like subsets.** A) Photo images of human intestinal gut and developing lymph nodes (arrows) in 8-17 PCW samples. Scale bar = 1cm. B) Bar plot with productive TCRαβ chain in fetal T and innate lymphoid cell types. C) Heatmap with scaled expression of selected differentially expressed genes in fetal LTi-like subsets, NK cells and adult ILC3 cells. D) UMAP visualisation and feature plots of Smart-seq2 processed flow cytometry-sorted CD45+ cells from second-trimester fetal tissue. UMAP visualisations are colored by intestinal region or leiden clustering. Feature plots show key gene expression described using 10X data. E) Bar graph showing the relative proportion (%) of cell types among total T and innate lymphocyte population across developmental and adult gut regions (FPIL-fetal proximal ileum, FMIL-fetal middle ileum, FTIL-fetal terminal ileum, FLI-fetal large intestine, FMLN-fetal mesenteric lymph node, DUO-duodenum, JEJ-jejunum, ILE-ileum, APD-appendix, CAE-caecum, ACL-ascending colon, TCL-transverse colon, DCL-descending colon, SCL-sigmoid colon, REC-rectum, MLN-mesenteric lymph node). F) UMAP visualisation with overlaid RNA velocity arrows of lymphocytes from fetal tissues colored by cell type. Zoom in panel shows ILCP and LTi-like ILC3 cells.

**Supplementary Fig. 6: Intestinal B cells and BCR analysis.** UMAP visualizations of B cells from fetal samples. CLP-common lymphoid progenitor B) Heatmap with mean expression of differentially expressed gene in fetal B cell populations as in A). C) Violin plot with MHCII score of fetal B cell subsets. UMAP visualisation colored by D) BCR isotype retrieved from 10X VDJ sequencing and E) 10x Genomics technology. BCR sequencing is available only for cells processed using 5’ 10x Genomics scRNA-seq technology. F) UMAP visualizations of B cells from pediatric and adult samples. LZ - light zone, DZ-dark zone, GC - germinal centre. G) Relative proportions (left) and count (right) of subtypes within total B cell factions in the small (above) and large (below) intestine separated by donor age (row). Units of age is years unless specified as weeks. H) Estimated clonal abundances per donor for members of expanded B cell clones in fetal and adult datasets. I) Quantitation of somatic hypermutation frequencies of IgH sequences from B cells in fetal and adult datasets. J) Quantitation of somatic hypermutation frequencies of IgH sequences (J) and Estimated clonal abundances per donor for members of expanded B cell clones (K) in gut regions (FPIL-fetal proximal ileum, FMIL-fetal middle ileum, FTIL-fetal terminal ileum, FLI-fetal large intestine, FMLN-fetal mesenteric lymph node, DUO-duodenum, JEJ-jejunum, ILE-ileum, APD-appendix, CAE-caecum, ACL-ascending colon, TCL-transverse colon, DCL-descending colon, SCL-sigmoid colon, REC-rectum, MLN-mesenteric lymph node). L) Binary count of co-occurrence of expanded B cell clones identified by single-cell VDJ analysis shared across gut regions and donors.

**Supplementary Fig. 7: Stromal and endothelial populations in the intestinal tract.** A) Heatmap with top differentially expressed genes between follicular dendritic cells (FDCs) and T reticular cells. B) Bar graph showing the relative proportion (%) of cell types among total stromal population across developmental and adult gut regions (FPIL-fetal proximal ileum, FMIL-fetal middle ileum, FTIL-fetal terminal ileum, FLI-fetal large intestine, FMLN-fetal mesenteric lymph node, DUO-duodenum, JEJ-jejunum, ILE-ileum, APD-appendix, CAE-caecum, ACL-ascending colon, TCL-transverse colon, DCL-descending colon, SCL-sigmoid colon, REC-rectum, MLN-mesenteric lymph node). C) Dendrogram plot of mesenchymal cells. D) Dotplot comparing key defining genes expressed across mLTo, FDC and T reticular cells. E) Heatmap of top differentially expressed genes between mLTo cells of different intestinal regions (FSI - fetal small intestine, FLI-fetal large intestine, FMLN-fetal mesenteric lymph nodes). UMAP projection of endothelial cell populations in fetal, pediatric and adult colored by annotation F) or cell cycle score G). Dashed line outlines lymphatic endothelial cell subsets (LEC). H) UMAP visualisation of subclustered LECs colored by cell cycle, subpopulation and sample age. I) Heatmap with top differentially expressed genes in the LEC subsets. J) Relative proportions (left) and count (right) of subtypes within total LEC factions separated by age (above) and intestinal region (below). Units of age is years unless specified as weeks. Region names are as in B). K) Feature plots with expression of lymphatic cells genes and key genes for lymphocyte adhesion. L) Violin plot of NFkB signalling activation score across LEC subpopulations.

**Supplementary Fig. 8: Ectopic lymphoid tissue formation in pediatric Crohn’s disease.** A) Heatmap with mean expression of ligand-receptor pairs in mLTo, LTi-like and LEC subsets as identified using CellphoneDB. B) Heatmaps with mean expression of curated immune recruitment signal genes in selected fetal stromal, epithelial and endothelial populations (top) and their receptor expression in the immune populations found in the fetal gut and mesenteric lymph nodes. C) UMAP visualisation of mesenchymal, smooth muscle and pericyte cell subsets in pediatric healthy and Crohn’s disease intestine colored by disease phenotype (left) and cell type annotation (right). mLTo-like cells are outlined using a dashed line. UMAP visualisation with overlaid RNA splicing and colored by pseudotime. D) Bar plots with number of cells in each donor grouped as control and Crohn’s disease (CD). E) Heatmap of genes reported in expanded mesenchymal cells in Rheumatoid arthritis (RA) or liver cirrhosis expressed by mLTo and mLTo-like cells in CD with enterocytes shown for reference.

**Supplementary Table1:** Sample metadata

**Supplementary Note Section 1:** Related to the cell type enrichment for IBD-GWAS genes analysis

## Materials and Methods

### Patient samples

Human fetal gut samples were obtained from the Human Developmental Biology Resource (HDBR, www.hdbr.org). The maternal consent was obtained through Newcastle hospital (the consent REC reference 18/NE/0290-IRAS project ID: 250012). Pediatric patient material used in intestinal organoid culture was obtained with informed consent as part of the ethically approved research study (REC-96/085).

Human adult tissue was obtained by the Cambridge Biorepository of Translational Medicine from deceased transplant organ donors after ethical approval (reference 15/EE/0152, East of England—Cambridge South Research Ethics Committee) and informed consent from the donor families. Fresh mucosal intestinal tissue and lymph nodes from the intestinal mesentery were excised within 1 hour of circulatory arrest; intestinal tissue was preserved in University of Wisconsin organ-preservation solution (Belzer UW Cold Storage Solution; Bridge to Life) and mLN were stored in saline at 4□°C until processing. Tissue dissociation was conducted within 2□ hours of tissue retrieval.

### Mouse samples

C57BL/6 mice were obtained from Jackson Laboratories (Margate, UK) and maintained in specific pathogen-free conditions at a Home Office-approved facility at the University of Cambridge. Female mice ages 10 to 14 weeks were used. Mice were housed in accordance with the United Kingdom Animals (Scientific Procedures) Act of 1986.

### Isolation of intestinal cells from fetal tissue

The fetal gut mesentery was cut to lengthen out the tissue and dissected into proximal ileum (PI), middle Ileum (PI), terminal Ileum (TI), colon and appendix. Samples were washed twice with Hanks Buffered Saline Solution (HBSS; Sigma-Aldrich; cat: 55021C) and minced into pieces using a scalpel. The samples were incubated in 2 ml HBSS solution containing 0.21 mg/ml Liberase TL (Roche; cat: 5401020001) or DH (Roche; cat: 5401089001) and 70 U/ml hyaluronidase (Merck; cat: 385931-25KU) for 50 minutes at 37°C shaking every 5 minutes and homogenised every 15 minutes using a pipette. The single cells were passed through a 40-100 μm sieve and spun down at 400 g at 4°C for 10 minutes. Red cell lysis solution (eBioscience™ 10X RBC Lysis Buffer (Multi-species) was used according to manufacturer’s guidelines to remove red blood cells, and the remaining cells were collected in FACS buffer (1% (v/v) FBS in PBS) by centrifugation at 400 g at 4°C for 5 minutes. All gut region samples (except mLN) proceed to MACS enrichment.

### Isolation of intestinal cells from adult tissue

Adult tissue sections were weighed before being washed in cold D-PBS (Gibco; cat: 14190094) and diced with a scalpel. Samples were dissociated in 1-2 ml of digestion mix (D-PBS, 250ug/ml Liberase TL (Roche; cat: 5401020001), 0.1 mg/ml DNaseI (Sigma; cat: 11284932001)) for 30 minutes at 37°C. Digested lysates were then passed through a 70 μm cell strainer, followed by 10 ml of neutralisation media (RPMI 1640 Medium with HEPES (Gibco; cat: 42401042), 20% (v/v) FBS (Sigma; cat: 25200-056)). The samples were then centrifuged at 700 g for 5 minutes at 4°C. Cells were resuspended in 1 ml 0.04% (w/v) BSA in D-PBS and counted using a NucleoCounter NC-200 and Via1-Cassette (ChemoMetec).

### MACS enrichment

Samples with greater than 500,000 cell yield were enriched by magnetic-activated cell sorting (MACS). Dissociated cells were centrifuged for 5 minutes at 300 g at 4°C and resuspended in 80 μl chilled MACS buffer (D-PBS, 0.5% (w/v) BSA (Sigma-Aldrich; cat: A7906-10G), 2 mM EDTA (ThermoFisher; cat: 15575020)) with 20 μl CD45 MicroBeads (Miltenyi Biotech; cat: 130-045-801) and incubated for 15 minutes at 4°C. Cells were washed with 2 ml MACS buffer and centrifuged as above and resuspended in 500 μl MACS buffer. Cells were passed through a pre-wetted MS column (Miltenyi Biotech; cat: 130-042-201) on a QuadroMACS Magnetic Cell Separator (Miltenyi Biotech) followed by four rounds of 500 μl of MACS buffer. Flowthrough was collected as the CD45-fraction. The column was removed from the magnet and the CD45+ fraction was eluted with 1 ml of MACS buffer. CD45- and CD45+ fractions were centrifuged as above and resuspended in 0.5-1 ml of 0.04% (w/v) BSA in D-PBS. Cell count and viability was determined using NucleoCounter NC-200 and Via1-Cassette (ChemoMetec) or hemocytometer and resuspended in 0.04% (w/v) BSA in D-PBS. Fetal CD45+ and CD45-fractions were combined at a 1:1 ratio.

### Organoid culture

Intestinal organoids from pediatric patients were cultured in Matrigel^®^ (Corning) using media described before. During organoid culture, the media was replaced every 48-72 hours. Organoids were passaged using mechanical disruption with a P1000 pipette and re-seeded in fresh growth-factor reduced Matrigel^®^ (Corning). When comparing culture media, multiple wells were seeded from a single dissociated sample. The organoids were then allowed to grow for 5 days followed by 24 hour treatment with recombinant human protein TNF-α (H8916, Sigma Aldrich) at 40ng/ml or IFN-gamma (PHC4031, Life Technologies Ltd) at 20ng/ml.

Processing for single-cell sequencing analysis was performed by removing the organoids from matrigel at passage 3-4 using incubation with Cell Recovery Solution at 4oC for 20 minutes, pelleting the cells, and re-suspending in TrypLE enzyme solution (Thermo Fisher) for incubation at 37°C for 10 mins. Cells were pelleted again and re-suspended in DMEM/F12.

### 10x Genomics Chromium GEX library preparation and sequencing

MACS enriched and total cell factions were loaded for droplet-based scRNA-seq according to the manufacturer’s protocol for the Chromium Single Cell 5’ gene expression v.2 (10x Genomics) to obtain 8,000-10,000 cells per reaction. Library preparation was carried out according to the manufacturer’s protocol. Pools of 16 libraries were sequenced across both lanes of an Illumina NovaSeq 6000 S2 flow cell with 50□ bp paired-end reads.

Intestinal organoids were prepared Chromium Single Cell 3’ gene expression v.2 (10x Genomics) to obtain 8,000 cells per reaction. Intestinal organoid cDNA libraries were sequenced on a single lane of an Illumina HiSeq 4000 with 50 bp paired-end reads.

### VDJ sample preparation

10x Genomics VDJ libraries were generated from the 5’ 10x Genomics Chromium complementary DNA (cDNA) libraries as detailed in the manufacturer’s protocol. BCR and TCR libraries for relevant samples were pooled and sequenced on a single lane of an Illumina HiSeq 4000 with 150 bp paired-end reads.

### Plate-based Smart-seq2

Plate-based scRNA-seq was performed with the NEBNext Single Cell/Low Input RNA Library Prep Kit for Illumina (catalog no. E6420L; New England Biolabs). Total cell fractions from dissociated gut sections of donors BRC2033-2034, F67, F72, F78 were snap-frozen in 10% (v/v) DMSO in 90% (v/v) BSA. Cells were thawed rapidly in a 37□ °C water bath and diluted slowly with a pre-warmed FACS buffer (2% (v/v) FBS in D-PBS). Cells were pelleted by centrifugation for 5□ min at 300 g, washed with 300□ μl of D-PBS and pelleted as before. Cells were resuspended in 100 μl of Zombie Aqua Fixable Viability Kit (1:200 dilution; catalog no. 423101) and incubated at room temperature for 15 minutes. Cells were washed with 2 ml of FACS buffer followed by 300 μl of FACS buffer and resuspended in a total of 100 μl of Brilliant Violet 650 mouse anti-human CD45 (Biolegend catalog no. 304043), Alexa Fluor 700 mouse anti-human CD4 (Biolegend, catalog no. 300526), and APC-H7 mouse anti-human CD19 (BD biosciences, catalog no. 560727) and incubated for 20□ minutes in the dark at room temperature. Cells were washed twice with 300□ μl of FACS buffer. Single, live, CD45+ cells were FACS sorted into wells of a 384-well plate (catalog no. 0030128508; Eppendorf) containing 2□ μl of 1× NEB Next Cell Lysis Buffer (New England Biolabs). FACS sorting was performed with a BD Influx sorter (BD Biosciences) with the indexing setting enabled. Plates were sealed and spun at 100 g for 1 min then immediately frozen on dry ice and stored at −80°C. cDNA and sequencing library generation was performed in an automated manner on the Bravo NGS Workstation (Agilent Technologies) as previously described ^6^. The purified pool was quantified on an Agilent Bioanalyzer (Agilent Technologies) and sequenced on one lane of an Illumina HiSeq 4000.

### Pre-processing of Smart-seq2 sequencing data

Cells with greater than 6000 genes and more than 25% mitochondrial reads were excluded, before regression of ‘n_counts’, ‘percent_mito’ and “G2M_score”. Cells positive for PTPRC expression (logTPM+1 >=0.2) were taken forward for downstream analysis.

### Pre-processing of 10x sequencing data

10x Genomics gene expression raw sequencing data were processed using the CellRanger software v3.0.0-v3.0.2 and the 10X human transcriptome GRCh38-3.0.0 as the reference. The 10x Genomics VDJ Ig heavy and light chains were processed using cellranger vdj v.3.1.0 and the reference cellranger-vdj-GRCh38-alts-ensembl-3.1.0 with default settings.

### scRNA-seq quality control and processing of 10x sequencing data

Pandas (version 0.24.2), NumPy (version 0.25.2), Anndata (version 0.6.19) and ScanPy (version 1.4) python packages were used to pool single-cell counts and for downstream analyses. Single-cell transcript counts for fetal and adult samples were handled separately to control for anticipated differences in cell expression and sample quality. Cells for each dataset were filtered for greater than 500 genes and less than 50% mitochondrial reads, and genes were filtered for expression in greater than 3 cells. Scrublet (version 0.2.1) score cuttoff of 0.25 was applied to assist in doublet exclusion. Additional doublet exclusion was performed throughout downstream processing based on unexpected co-expression of canonical markers such as CD3D (component of the TCR) and EPCAM. Gene expression for each cell was normalised and log transformed. Cell cycle score was calculated using expression of 97 cell cycle genes listed in Tirosh et al. 2015 ^70^. Cell cycle genes were then removed for initial clustering. Cell cycle score, mitochondrial reads percentage and unique molecular identifiers (UMIs) were regressed prior to scaling the data.

### Cell type annotation and scoring

Batch correction of fetal and adult datasets were performed with bbknn (version 1.3.9, neighbours=2-3, metric=‘euclidean’, n_pcs=30-50, batch_key = “fetal_id” or “batch”), and dimensionality reduction and leiden clustering (resolution 0.3-1.5) was carried out and cell lineages were annotated based on marker gene expression for each cluster. Cell lineages were then subclustered and cell cycle genes reintroduced for batch correction and leiden clustering for annotation of cell types and states again based on marker gene expression. Annotated fetal and adult datasets were then merged and annotations adjusted for concordance.

For MHCII scoring of M cells and B cells we use the following genes: *SECTM1, CD320, CD3EAP, CD177, CD74, CIITA, RELB, TAP2, HLA-DRA, HLA-DRB5, HLA-DRB1, HLA-DQA1, HLA-DQB1, HLA-DQB1-AS1, HLA-DQA2, HLA-DQB2, HLA-DOB, HLA-DMB, HLA-DMA, HLA-DOA, HLA-DPA1, HLA-DPB1*. Glial or neuronal cell signature score was calculated using the curated gene sets (glial: *SOX10, MPZ, S100B, ERBB3, PLP1, GAS7, COL18A1, EDNRB;* neuronal: *ELAVL3, ELAVL4, TUBB2B, PHOX2B, RET, CHRNA3*).

The scoring was done using sc.tl.score_genes() function with default parameters to calculate the average expression of selected genes substrated with the average expression of reference genes.

### Intestinal organoid analysis

Single-cell count matrices from three organoid growth conditions were combined together using Pandas (version 0.24.2) and NumPy (version 0.25.2) packages. Cells with fewer than 8000 genes and with less than 20% mitochondrial reads were included in the analysis. Genes with expression in fewer than 3 cells were also excluded. For *PLCG2* expression comparison we use normalised (sc.pp.normalize_per_cell) and log transformed (sc.pp.log1p) counts. The data was plotted using Seaborn package barplot and swarmplot functions (version 0.11.0).

### Differential cell-state abundance analysis for BEST4+ cells

To identify region-specific subpopulations, we performed compositional analysis between *BEST4+* enterocytes from small and large intestine tissue, using an unpublished tool for differential abundance testing on KNN graph neighbourhoods (implemented in the R package miloR https://github.com/MarioniLab/miloR).

Briefly, we performed PCA dimensionality reduction and KNN graph embedding on the *BEST4*+ enterocytes. We define a neighbourhood as the group of cells connected to a sampled cell by an edge in the KNN graph. Cells are sampled for neighbourhood construction using the algorithm proposed previously ^71^. For each neighbourhood we then perform hypothesis testing between conditions to identify differentially abundant cell states whilst controlling the FDR across the graph neighbourhoods.

We test for differences in abundance between the cells from small and large intestine tissue in adult samples and fetal samples. To identify markers of SI-specific and LI-specific subpopulations, we performed differential gene expression analysis using a linear model implemented in the package limma ^72^, using 1% FDR (implemented in the function findNhoodMarkers of the miloR package).

### RNA velocity, monocle 3 and diffusion map pseudotime analyses

For neuronal cell trajectory analysis we use scVelo 0.21 package implementation in Scanpy 1.5.1 ^73^. The data was sub-clustered on fetal neuronal cells and pre-processed using functions for detection of minimum number of counts, filtering and normalisation using scv.pp.filter_and_normalise and followed by scv.pp.moments function. The gene-specific velocities were obtained using scv.tl.velocity(mode=‘stochastic’) and scv.tl.velocity_graph() by fitting a ratio between unspliced and spliced mRNA abundances. The gene specific velocities were visualised using scv.pl.velocity_graph() or scv.pl.velocity_embedding_grid() functions.

In addition, we calculate pseudotime using Monocle 3 0.2.1 (R version 3.6.3) ^74–77^In short, we use preprocess_cds(num_dim=100) to normalise and pre-process raw counts, and reduce dimension and cluster cells using functions reduce_dimension(reduction_method = ‘UMAP’) and cluster_cells(). We then use functions learn_graph() and order_cells (root cell = “GTATTCTGTCATGCCG-1-4918STDY7426908”) to order cells along the pseudotime. We then overlay the pseudotime onto the UMAP with velocyto arrows using scv.pl.velocity_embedding_grid() with clusters set to monocle3 inferred “pseudotime”. To visualise genes that change along the pseudotime we use sc.pl.paga_path() function with pseudotime set to monocle3 pseudotime. This function required calculation of PAGA parameters and dpt_pseudotime with functions as follows: sc.tl.paga(), sc.pl.paga(), sc.tl.draw_graph (init_pos=‘paga’), sc.tl.dpt() ^78^.

### BCR analysis

Single-cell BCR analyses were performed as described previously ^79^. Briefly, poor quality or incomplete VDJ contig sequences were discarded and all IgH sequences for each donor were combined together. IgH sequences were annotated with IgBLAST ^80^ before isotype reassignment using AssignGenes.py (pRESTO; ^81^). Ambiguous V gene calls were corrected using TIgGER v.03.1 ^82^ before identifying clonally-related sequences with DefineClones.py (ChangeO v.0.4.5; ^82^) using a threshold of 0.2 for nearest neighbour distances. The germline IgH sequence for each clonal family was determined using CreateGermlines.py (ChangeO v.0.4.5) followed by using observedMutations (Shazam v.0.1.11; ^82^) to calculate somatic hypermutation frequencies for individual sequences. Finally, for integration with the singlecell gene expression object, the number of high quality and annotated contigs per Ig chain (IgH, IgK, IgL) was determined for each cell barcode. If multiple unique sequences for a given chain were detected, that cell was annotated as “Multi” and not considered in further analysis. BCR metadata was combined with the scRNA object for downstream analysis and comparison of different B cell populations.

### Cell-cell communication analysis

To infer cell-cell communication and screen for ligand-receptors involved we applied CellPhoneDB v2.0 python package ^83,84^ on the normalised raw counts and fine cell type annotations from the second trimester intestinal samples (12-17 PCW). We use default parameters and set subsetting to 5000 cells. To identify most relevant interactions, we subset specific interactions based on the ligand/receptor expression in more than 10% of cells within a cluster and where the log2 mean expression of the pair is greater than 0.

### Enteric nervous system disease-linked gene expression

Hirschsprung’s disease genes were curated from ^35–37^ and The Human Phenotype Ontology website (see URL: https://hpo.jax.org/app/, Aganglionic megacolon HP:0002251) and we selected genes with higher than or equal to 0.3 expression in neuronal lineage single cells. We calculated mean expression per cluster and organ and visualised the normalised mean expression (standard_scale=“var”) using seaborn.clustermap() function (version 0.11.0).

### IBD-GWAS gene expression analysis in epithelial cells in Figure 2D

Crohn’s disease (EFO_0000384) and ulcerative colitis (EFO_0000729) gene names were retrieved from GWAS Catalog of EMBL-EBI (https://www.ebi.ac.uk/gwas/). The list was downloaded on 27 May 2020. We took the intersecting genes from pooled GWAS genes (P value < 1.00E-6) and the top 100 positive and 100 negative differentially expressed marker genes for each epithelial cell type versus all other epithelial cell types. Expression colour scale has a maximum cut off of 1.5 mean normalised counts to increase resolution at lower expression levels. Only cell types present are shown in each respective gut region heatmap.

### Cell type enrichment for IBD-GWAS genes

IBD GWAS summary statistics of Crohn’s disease (CD) and ulcerative colitis (UC) were obtained from the International Inflammatory Bowel Disease Genetics Consortium (IIBDGC) (see URL: https://www.ibdgenetics.org/). The GWAS enrichment analysis of 103 annotated gut cell types for CD and UC was performed using a fGWAS approach ^85,86^, often used for fine-mapping and enrichment analysis of various functional annotations for molecular quantitative trait and GWAS loci. The association statistics (log odds ratios and standard errors) were converted into the approximate Bayes factors using the Wakefield approach ^87^. A *cis*-regulatory region of 1Mb centred at the transcription start site (TSS) was defined for each gene (Ensembl GRCh37 Release 101). The Bayes factors of variants existing in each *cis* region were weighted and averaged by the prior probability (an exponential function of TSS proximity) estimated from the distance distribution of regulatory interactions ^88^. The likelihood of an fGWAS model was given by the averaged Bayes factors across all genome-wide genes multiplied by the feature-level prior probability obtained from a linear combination of cell type specific expression and the averaged expression across all cell types as a baseline expression. The enrichment of each cell type was estimated as the maximum likelihood estimator of the effect size for the cell type specific expression. The code of the pairwise hierarchical model (see URL: https://github.com/natsuhiko/PHM) was utilised for the enrichment analysis. The detailed model derivation is demonstrated in the Supplementary Note Section 1.

### Cryosectioning, smFISH, and confocal imaging

Fetal gut tissue was embedded in optimal cutting temperature medium (OCT) and frozen on an isopentane-dry ice slurry at −60°C, and then cryosectioned onto SuperFrost Plus slides at a thickness of 10 μm. Prior to staining, tissue sections were post-fixed in 4% paraformaldehyde in PBS for 15 minutes at 4°C, then dehydrated through a series of 50%, 70%, 100%, and 100% ethanol, for 5 minutes each. Staining with the RNAscope Multiplex Fluorescent Reagent Kit v2 Assay (Advanced Cell Diagnostics, Bio-Techne) was automated using a Leica BOND RX, according to the manufacturers’ instructions. Following manual pretreatment, automated processing included epitope retrieval by protease digestion with Protease IV for 30 minutes prior to RNAscope probe hybridisation and channel development with Opal 520, Opal 570, and Opal 650 dyes (Akoya Biosciences). Stained sections were imaged with a Perkin Elmer Opera Phenix High-Content Screening System, in confocal mode with 1 μm z-step size, using a 20× water-immersion objective (NA 0.16, 0.299 μm/pixel). Channels: DAPI (excitation 375 nm, emission 435-480 nm), Opal 520 (ex. 488 nm, em. 500-550 nm), Opal 570 (ex. 561 nm, em. 570-630 nm), Opal 650 (ex. 640 nm, em. 650-760 nm).

### Flow cytometry validation of Fcgr on mice tuft cells

Small intestines from C57BL/6 mice were flushed of faecal content with ice-cold PBS, opened longitudinally, cut into 0.5 cm pieces, and washed by vortexing three times with PBS with 10mM HEPES. Tissue was then incubated with an epithelial stripping solution (RPMI-1640 with 2% (v/v) FCS, 10mM HEPES, 1mM DTT, and 5 mM EDTA) at 37°C for two intervals of 20 minutes to remove epithelial cells. The epithelial fraction was subsequently incubated at 37°C for 10 minutes with dispase (0.3 U/mL, Sigma-Aldrich) and passed through a 100μm filter to obtain a single-cell suspension. Cells were blocked for 20 minutes at 4°C with 0.5% (v/v) heat-inactivated mouse serum followed by extracellular staining in PBS at 4°C for 45 minutes with the following antibodies; EpCAM-FITC (1:400, G8.8, Invitrogen), CD45-Bv650 (1:200, 30-F11, BioLegend), CD11b-Bv421 (1:300, M1/70, BD Biosciences), Siglec-F-APC (1:200, 1RNM44N, Invitrogen), FcγRI-PE (1:200, X54-5/7.1, BioLegend), FcγRIIB-PE (1:200, AT130-2, Invitrogen), FcγRII/RIII-PE (1:200, 2.4G2, BD Biosciences), FcγRIV-PE (1:200, 9E9, BioLegend) and Rat IgG2b, κ isotype-PE-Cy7 (1:200, LOU/C, BD Biosciences). Cells were then stained with LIVE/DEAD Fixable Aqua Dead Cell Stain Kit (Thermo Fisher Scientific) for 20 minutes at room temperature, fixed with 2% PFA, and analysed on a CytoFLEX LX (Beckman Coulter) flow cytometer.

### Data availability

The expression data for fetal and adult regions is available in an interactive browsing website: www.gutcellatlas.org. Raw sequencing data are available at ArrayExpress (accession numbers E-MTAB-9543, E-MTAB-9536, E-MTAB-9532 and E-MTAB-9533).

### Code availability

Processed single-cell RNA sequencing objects will be available for online visualisation and download at gutcellatlas.org. The code generated during this study will be available at Github https://github.com/teichlab/.

