## Supplementary figures and images for "Cells of the human intestinal tract mapped across space and time"

### Supplementary figures

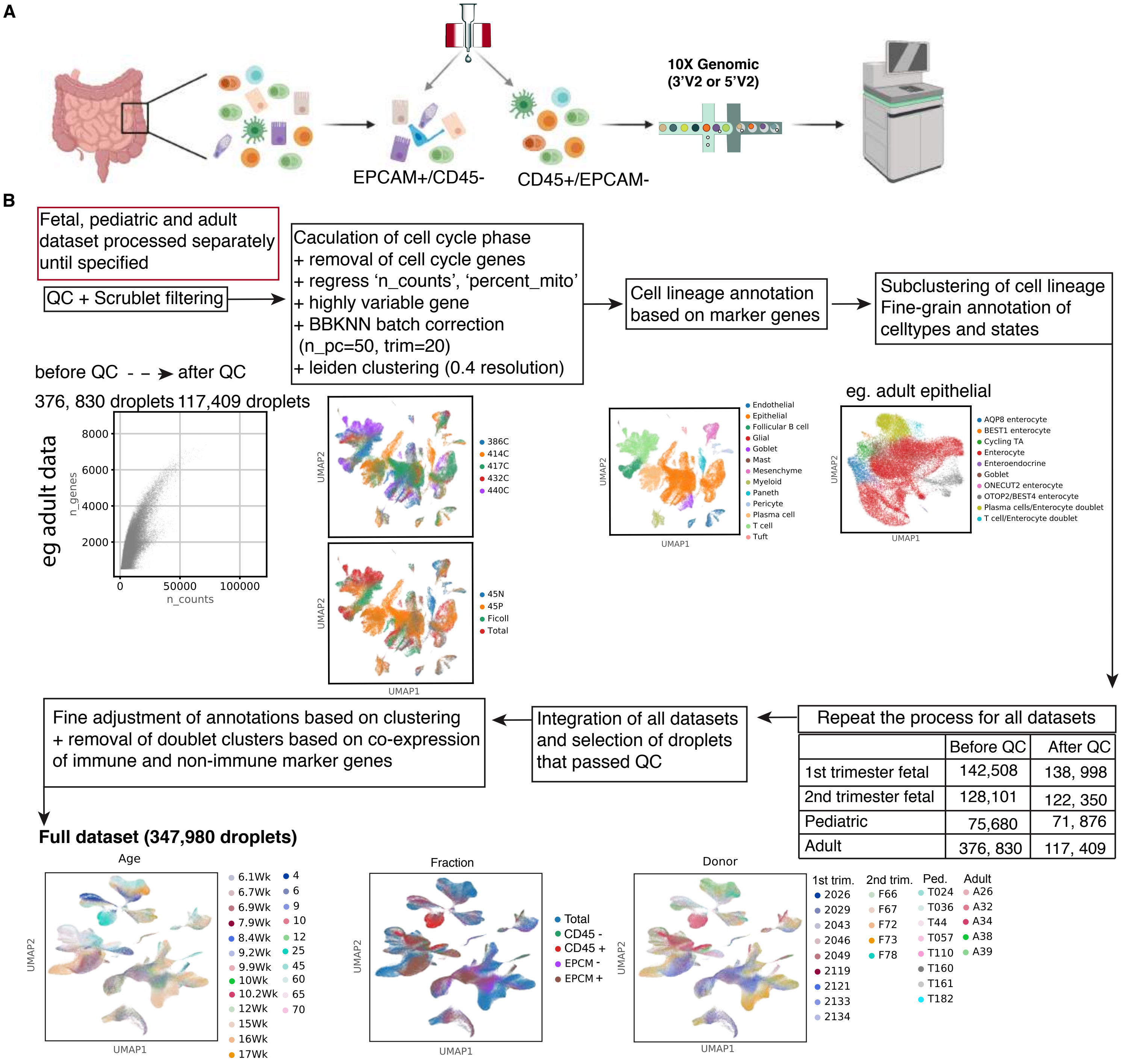


| | Before QC | After QC |
|---|---|---|
| 1st trimester fetal | 142,508 | 138, 998 |
| 2nd trimester fetal | 128,101 | 122, 350 |
| Pediatric | 75,680 | 71, 876 |
| Adult | 376, 830 | 117, 409 |

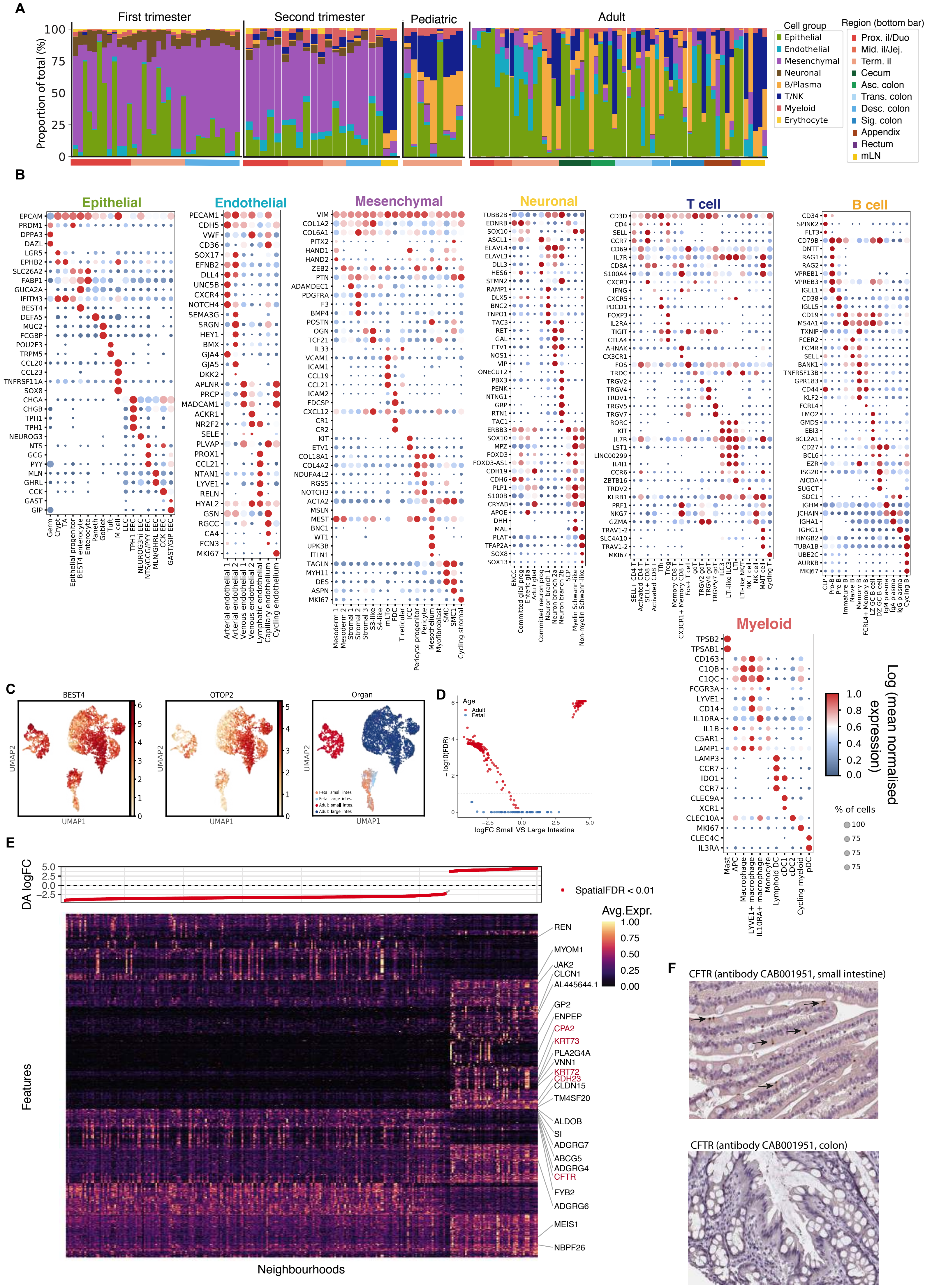

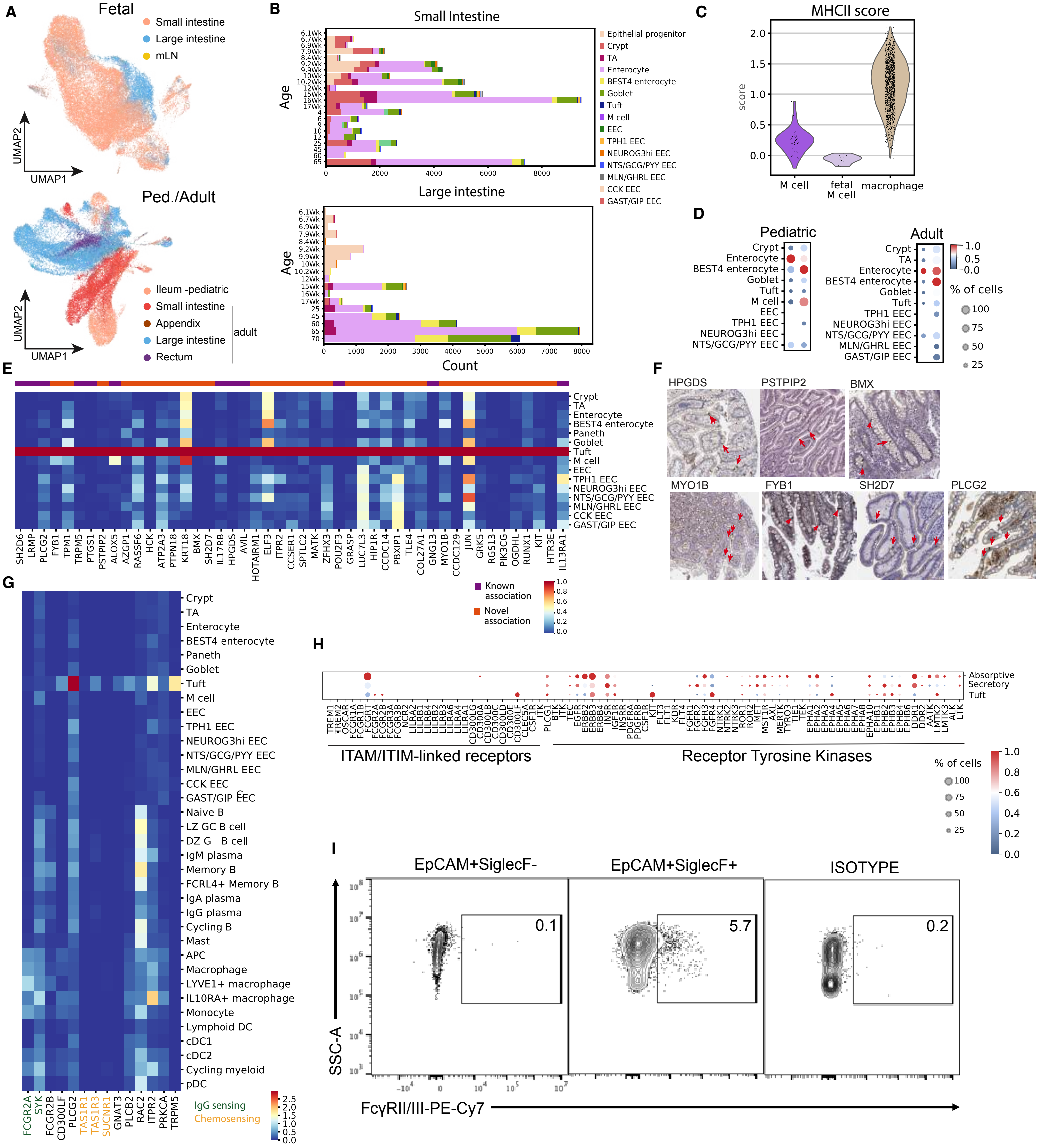

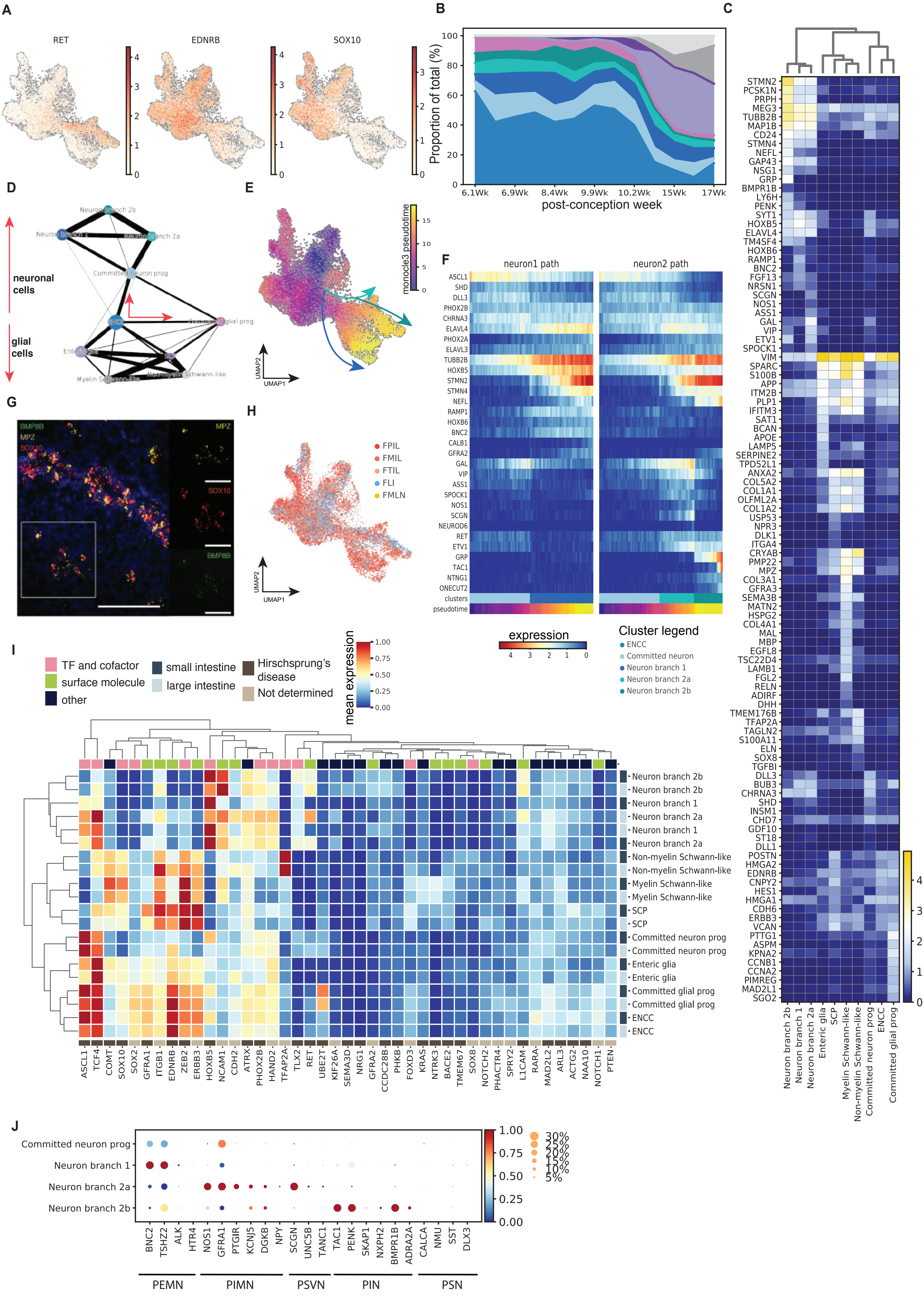

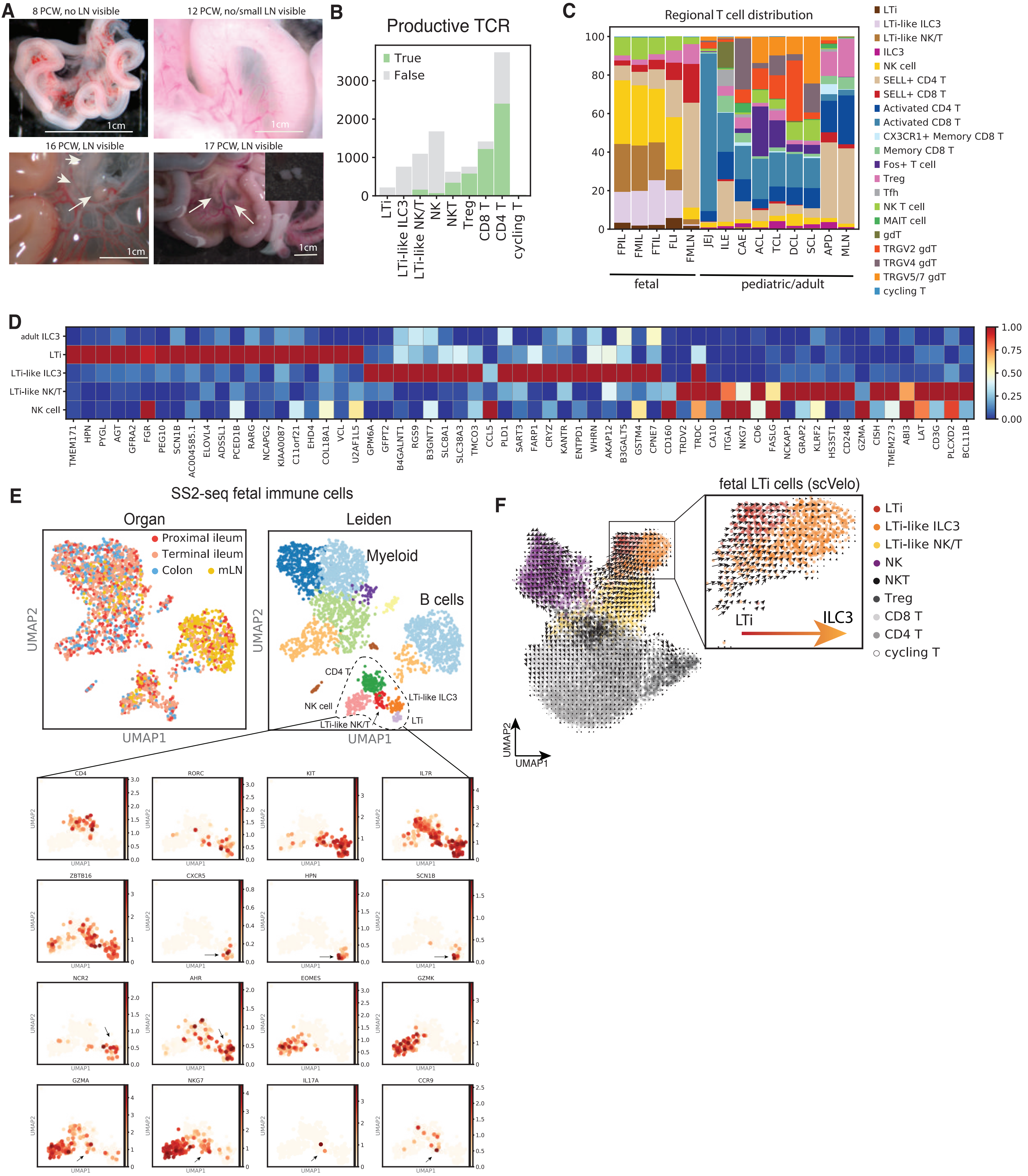

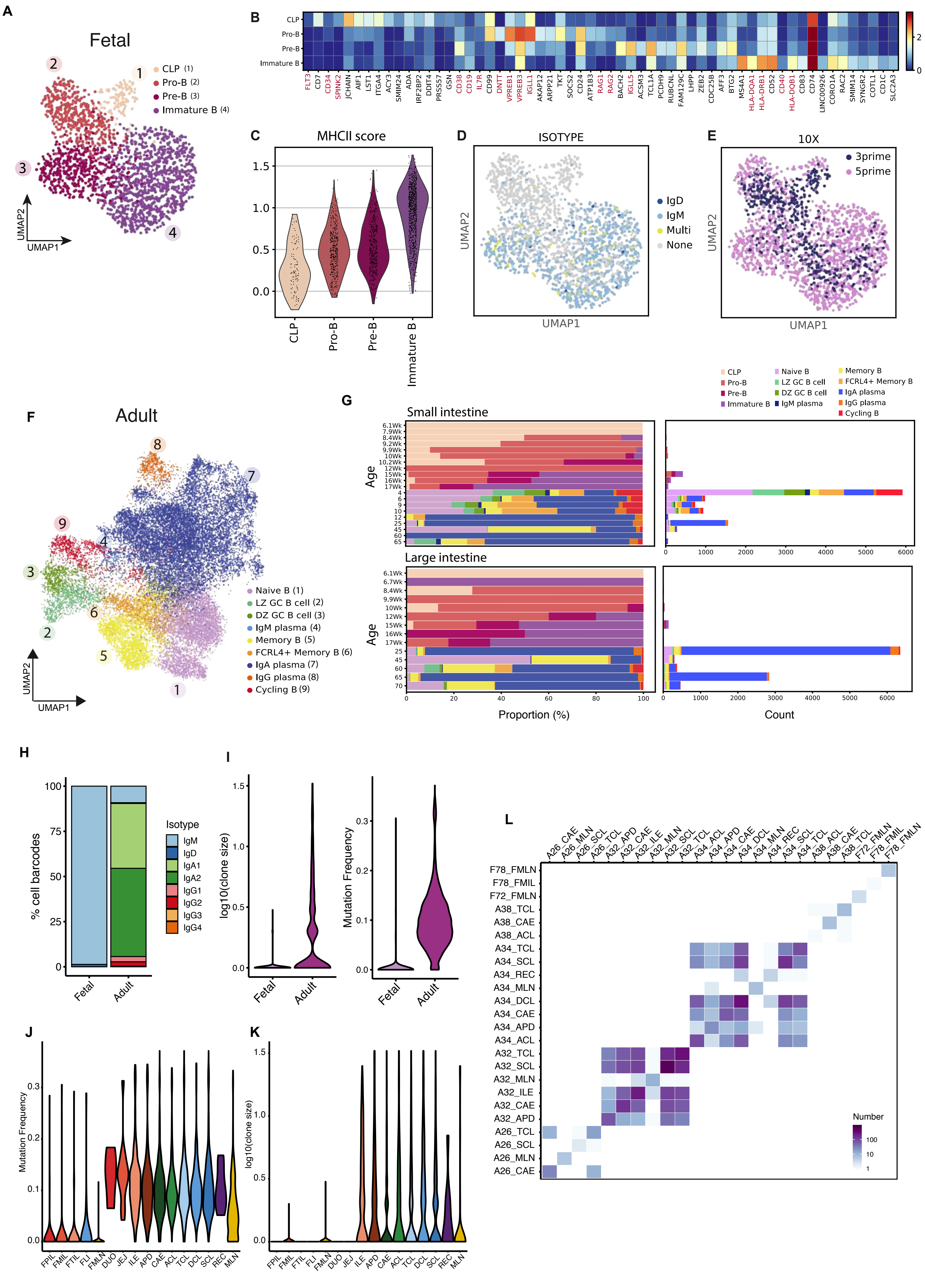

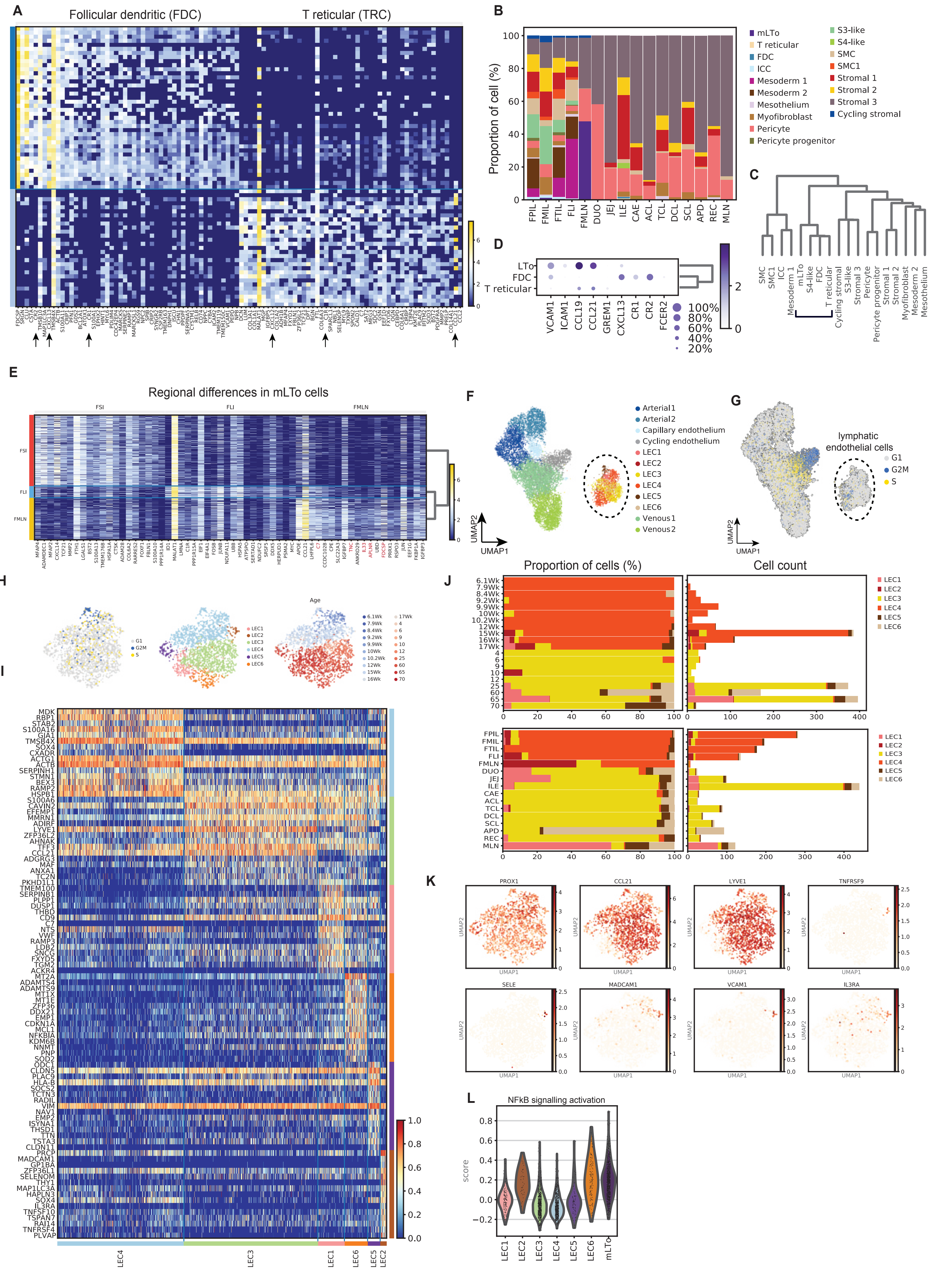

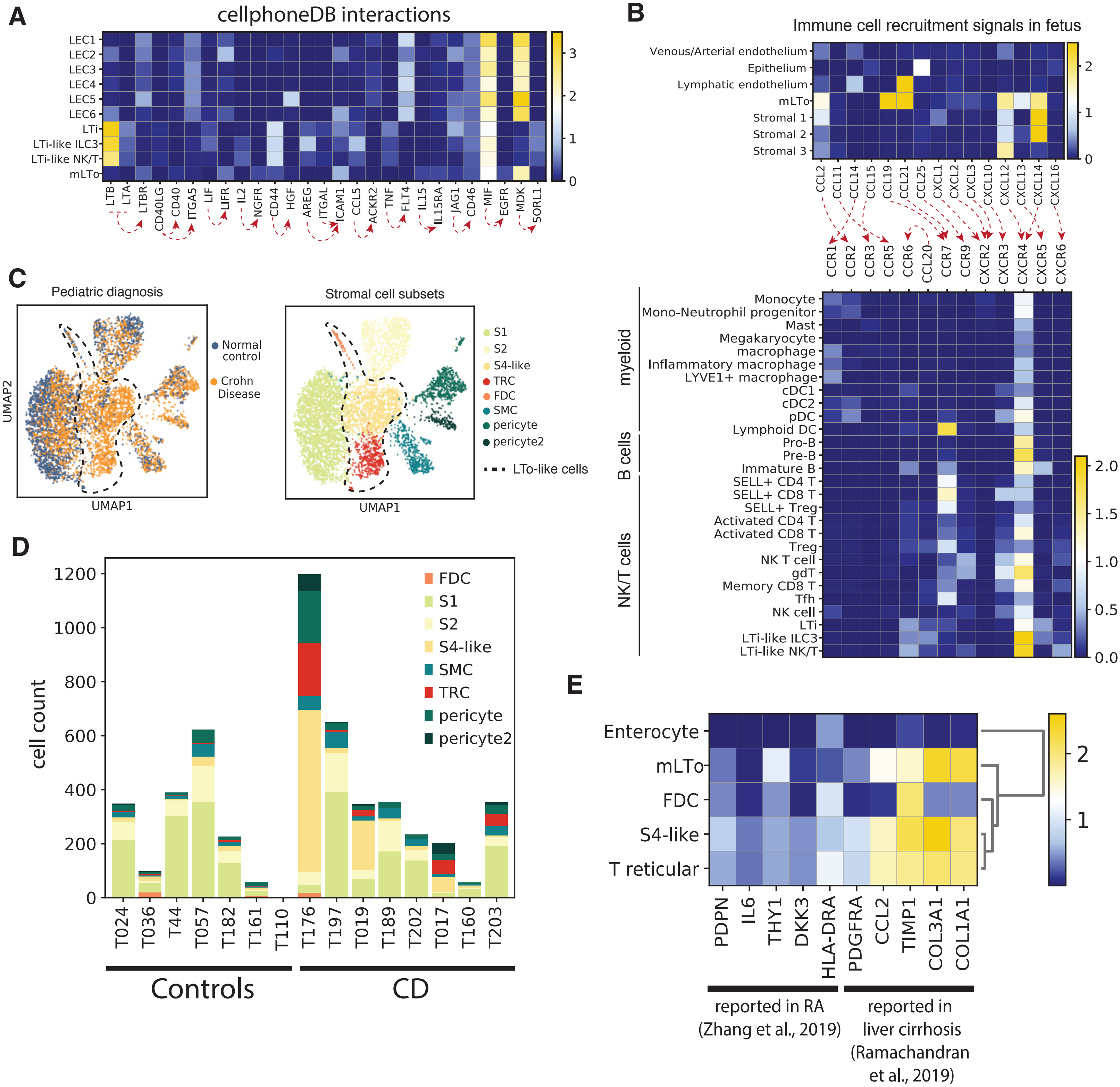
